# The complete cell atlas of an aging multicellular organism

**DOI:** 10.1101/2022.06.15.496201

**Authors:** Antoine E. Roux, Han Yuan, Katie Podshivalova, David Hendrickson, Rex Kerr, Cynthia Kenyon, David R. Kelley

**Affiliations:** Calico Life Sciences LLC, South San Francisco, California 94080, USA

**Author notes:** Authors contributed equally (Antoine E. Roux, Han Yuan).

## Abstract

Here we describe a single-cell atlas of aging for the nematode *Caenorhabditis elegans.* This unique resource describes the expression across adulthood of over 20,000 genes among 211 groups of cells that correspond to virtually every cell type in this organism. Our findings suggest that *C. elegans* aging is not random and stochastic in nature, but rather characterized by coordinated changes in functionally related metabolic and stress-response genes in a highly cell-type specific fashion. Aging signatures of different cell types are largely different from one another, downregulation of energy metabolism being the only nearly universal change. Some biological pathways, such as genes associated with translation, DNA repair and the ER unfolded protein response, exhibited strong (in some cases opposite) changes in subsets of cell types, but many more were limited to a single cell type. Similarly, the rates at which cells aged, measured as genome-wide expression changes, differed between cell types; some of these differences were tested and validated *in vivo* by measuring age-dependent changes in mitochondrial morphology. In some, but not all, cell types, aging was characterized by an increase in cell-to-cell variance. Finally, we identified a set of transcription factors whose activities changed coordinately across many cell types with age. This set was strongly enriched for stress-resistance TFs known to influence the rate of aging. We tested other members of this set, and discovered that some, such as GEI-3, likely also regulate the rate of aging. Our dataset can be accessed and queried at c.elegans.aging.atlas.research.calicolabs.com/.

## Introduction

The nematode *Caenorhabditis elegans* is a powerful model organism for studying multicellular development and lifespan. Identification of genes controlling lifespan in *C. elegans* has allowed the discovery of cellular regulators of aging conserved in other organisms, including mammals (Kenyon, 2010). However, our understanding of these mechanisms at the cellular and molecular levels remains incomplete, in part because genome-wide analyses of *C. elegans* have been mostly limited to whole-animal studies, with isolated tissues rarely profiled and compared. Thus, the research community lacks data describing aging at the cellular level, in every tissue, and during the entire course of adulthood.

Recent advances in transcriptomics have enabled scientists to profile gene expression at the level of single cells. The biological accuracy of single cell RNA sequencing (scRNA-seq) surpasses that of bulk RNA-seq because it does not average gene expression across the entire cell population, organ or organism (Hwang et al., 2018). Studies comparing the effect of aging on single-cell gene expression in mice have revealed tissue-specific aging signatures, as well as shared aging genes regulated similarly between cell types (Kimmel et al., 2019; Tabula Muris Consortium, 2020; Zhang et al., 2021). However, due to the large number of cells and organ complexity in mice, these datasets remain incomplete, omitting many organs (for example several muscles, peripheral blood, gonads, esophagus, stomach, tail and more than half of the brain) and many connective tissues (Tabula Muris Consortium et al., 2018). A complete view of an entire multicellular organism’s aging is not yet available.

Application of scRNA-seq to *C. elegans* is especially opportune for several reasons. *C. elegans* contains a small, defined number of cells (959 somatic cells). Cellular lineages and physical locations are comprehensively described for each cell of the adult animal. The animal is transparent, and knowledge accumulated from gene tagging *in vivo* is substantial, providing many marker genes to help identify cell types. *C. elegans*’ short lifespan and its experimental and genetic tractability greatly facilitates follow up studies. Recently, scRNA-seq, in combination with a cell-dissociation protocol (Zhang et al., 2011), has been applied successfully to study *C. elegans* embryogenesis and larval development (Cao et al., 2017; Packer et al., 2019; Taylor et al., 2021; Tintori et al., 2016). However, comprehensive examination of adult animals using this approach has been hindered by the abundance of germ cells, which comprise roughly two thirds of the adult cells (Kimble and Hirsh, 1979), as well as a bias toward germ-cell isolation during dissociation (Cao et al., 2017; Packer et al., 2019) (personal communication and this study).

In this study, we developed experimental and bioinformatic methods for depleting germ cells that enabled the collection of gene expression data in nearly every somatic cell type across the adult lifespan of *C. elegans*. We identified 211 unique gene expression profiles and matched the large majority to known cell types. We verified the quality of our cell-identity annotation and novel marker genes using microscopy and comparison to published pre-adult scRNA-seq datasets. We then asked which segments of the aging transcriptome signatures were cell-type specific and which were shared. Our data revealed a high level of tissue-specific changes in gene expression, including activation of processes that might restore homeostasis, such as the up-regulation of proteostasis and DNA-repair genes in certain tissues. The shared signature of aging includes decreased expression of metabolic genes. We measured the extent to which the transcriptome changed during aging and found that the magnitude was much larger for some cell types than for others. We validated this inferred difference in cell aging rates *in vivo* by examining age-dependent changes in mitochondrial morphology. As expected, the transcriptional signatures of some tissues became more noisy with age; however, unexpectedly, the signatures of others became more cohesive. We quantified the expression and inferred activities of mRNAs encoding over 200 transcription factors (TFs) in every tissue throughout adulthood. We made the unexpected discovery that many transcription factors that can extend lifespan are naturally up-regulated with age, and based on these TF expression signatures, we identified new TFs that appear to regulate aging. Finally, we developed an online public interface to make genes, gene sets and inferred TF activity easy to explore in every cell type at every age [c.elegans.aging.atlas.research.calicolabs.com/].

## Results

### Single-cell RNA sequencing of aging *C. elegans*

We set out to perform time series scRNA-seq on a *C. elegans* population over the course of their adult lifespan and for that purpose selected six, roughly evenly spaced, time points (days 1, 3, 5, 8, 11 and 15 of adulthood, Figure 1A). We used three methods to reduce germ cell number. First, we analyzed temperature-sensitive *gon-*2*(q388ts)* mutants, which fail to develop gonads at the non-permissive temperature (Sun and Lambie, 1997). Due to the incomplete penetrance of this mutation, we were still able to detect gonad dependent gene expression cell signatures, and we still observed an overrepresentation of germ cells in the data, as determined by germline marker gene set enrichment scores (Suppl. Notes 1) (Aibar et al., 2017). To further enrich somatic cells, we FACS sorted cells based on their genetic ploidy using DAPI, as the majority of the germ cells are meiotically arrested with a 4N ploidy. This method greatly enriched somatic cells but still did not eliminate the germ-cell signal altogether. Therefore, we computationally identified and removed any remaining germ cells, as well as embryonic cells and sperm cells, from the dataset (see Methods). Together, our methods enriched somatic cells up to a factor of >7 times at the older time points.

**Figure 1.**
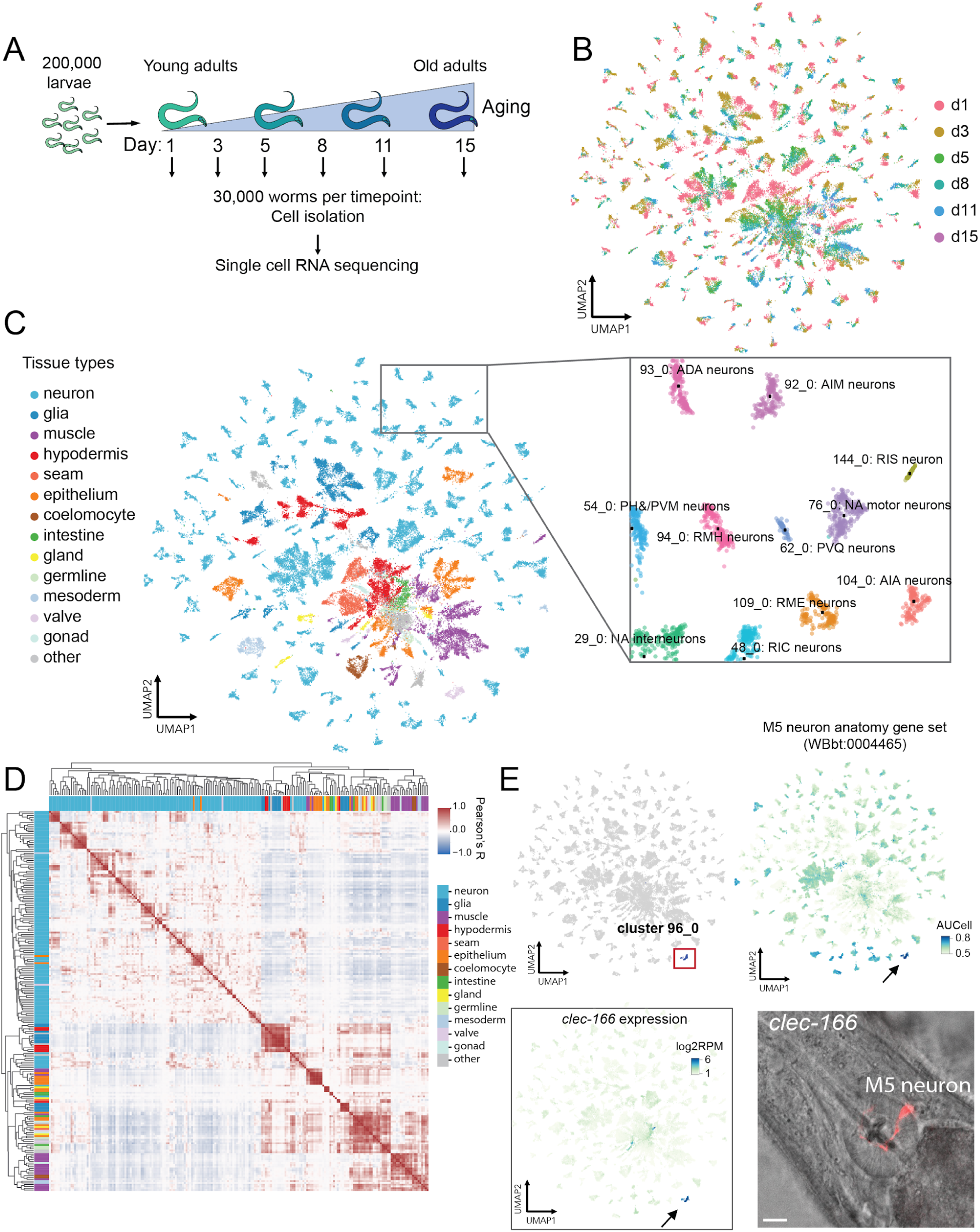
Single-cell RNA sequencing of C. elegans adults revealed 211 clusters covering nearly every known cell type at six ages. A) Experimental design. B) UMAP representation of all cells colored by time point. C) UMAP representation of all cells colored by high-level cell type annotation (tissue types, detailed in Table S1). Inset shows specific annotations for several neuron clusters. For UMAP visualization of all 211 clusters, see Figure S1D. D) Heatmap of Pearson correlations between every pair of cell-type clusters’ average gene expression. High-level tissue type is annotated in the color bar. This matrix is annotated in Figure S1F. E) Example of a new marker gene. Top left: cells belonging to cluster 96_0 are labeled in blue; Top right: UMAP coloring cells by AUCell score of WBbt:0004465, anatomy term gene set for M5 neuron; Bottom left: UMAP coloring cells by expression of our newly-identified M5 marker gene, clec-166. Bottom right: Visualization of M5 by promoter_clec-166::scarlet expression. Arrows point to cluster 96_0. Scale bar: 10𝝻m.

We applied CellBender (Fleming et al., 2019) to remove empty droplets and correct for ambient background RNA. We performed additional cell filtering in cases where CellBender ambient RNA removal was incomplete (see Methods). The final dataset contained 47,423 cells quantifying 20,305 genes across the full time series. Our method yielded more reads per cell than previous scRNA-seq in *C. elegans* adults (2,175 unique molecular identifiers (UMIs) vs. 156 UMIs per cell, 644 genes vs. 52 genes per cell, comparing our data to that of Preston et al. (Preston et al., 2019). However, we note that at the last time point (d15), we recovered fewer cells and fewer UMIs per cell (Figure S1A-C).

We embedded the cells into a low-dimensional latent space using the negative binomial variational autoencoder scVI and computed a nearest neighbor graph (Lopez et al., 2018). Uniform Manifold Approximation and Projection (UMAP) visualization (McInnes et al., 2018) revealed considerable cluster structure and variation with age (Figure 1B, S1D).

### Cell type annotation

Most adult *C. elegans* cell types lack comprehensive gene expression profiles, but many have sparse marker-gene data collected from published microscopy studies. To annotate cell types *de novo* without age as a confounder, we encoded and decoded the raw gene expression counts through scVI, treating time points as batches, to form denoised and age-corrected profiles. We then applied Leiden community detection to the nearest neighbor graph and identified 147 super-clusters of cells (Traag et al., 2019). Noticing clear substructure within super-clusters, we further applied Leiden to each super-cluster individually, tuning for optimal resolution using functionally relevant metrics to identify a total of 211 cell clusters (Figure S1D). All time points except for d15 were represented in most clusters (Figure S1A), confirming that the clusters capture cell type variation rather than age variation (Figure S1C, S1E). Nevertheless, visualizing each cluster on the original UMAP without any age correction, we observed a clear continuous aging trajectory within most clusters (Figure S1E).

To annotate these cell clusters with known cell-type identities, we queried the 211 clusters using 532 WormBase anatomy gene sets using three different approaches. Here, and in the analyses to follow, we leveraged a statistic called AUCell, which quantifies gene set enrichment at the top of a ranked gene list, where the genes are ranked by expression in a single cell (Aibar et al., 2017). We used an age-corrected scVI model to perform clustering and annotation to avoid confounding cell type and cell changes with age.

In our first approach, we applied AUCell to individual cells by computing enrichment scores for each WormBase anatomy gene set, summarizing these scores at the cluster level and assigning each anatomy term to its top three clusters.

For our second approach, we established a set of marker genes for each cluster using differential expression compared to all cells outside of the cluster. We then quantified the enrichment of each anatomy gene set with a hypergeometric test for significant overlap. In our final approach, we manually visualized the AUCell score distribution of each WormBase anatomy gene set on the UMAP, assigning the anatomy term to one or multiple clusters. The three annotation methods usually resulted in consistent anatomy annotation, but the AUCell method tended to bias toward smaller anatomy terms and the hypergeometric test tended to bias toward larger anatomy terms (Table S1). We summarized the results from the three methods and assigned a final annotation to each cluster (Table S1).

Together the clusters comprised a large portion of the known cell types from every tissue, including the intestine, hypodermis, seam cells, various muscles, glands, glia and neurons (Figure 1C, Table S1). 73% of the clusters were described by a single anatomy term (cell type), 18% by more than one anatomy term. 9% did not enrich for any anatomy term, which we call ‘orphan’ clusters. For many of these orphan clusters, assignment was hampered by the lack of known marker genes. Improving these annotations will be an ongoing process as more single-cell and *in vivo* expression data are collected. We assigned putative annotations to orphan clusters as detailed below.

To further the analysis, we assigned a high-level annotation (tissue type) to each specific anatomy term (Figure 1C). Neurons comprised the most abundant cell type, with 133 distinct clusters. Muscles were also represented by multiple cell types, 22 in all. Because our germ cell removal remained incomplete, we also identified several germline clusters. Due to the partial penetrance of the *gon-2(q388)* mutation, our cell population also contained somatic gonads. Importantly, comparing these clusters with those of a strain carrying a wild-type copy of *gon-2*, we showed that the mutation did not alter the expression of genes assigned to gonad cells (Suppl. Notes 1), and we recovered non-gonadal cells whose development requires signals from the gonad (for example, vulval cells, (Ferguson and Horvitz, 1985). Finally, we note that since our germline removal approach might have introduced selection bias for germ cell subtypes (as 4N cells were removed), these clusters must be interpreted accordingly.

To visualize the clusters from the perspectives of their annotations, we compared clusters with each other using a pairwise Pearson’s correlation of the average cluster expression. This revealed a prominent bifurcation separating neurons and non-neuronal cells (Figure 1D, S1F). Zooming in, both the neuronal and non-neuronal groups exhibited considerable structure arising from functionally similar and different cell types. Multiple clusters annotated as glia, muscles and coelomocytes grouped together in this correlation matrix, reflecting their similar gene expression. Based on the similarities of different cell types revealed by hierarchical clustering, we were able to assign candidate annotations to the orphan clusters (Figure S1F, Table S1).

These cell-type similarities suggested new insights into their functions (Figure S1F), particularly for tissues that had never been isolated before. For example, the excretory canal CAN neuron (69_0) was not clustered among the neuronal clusters but rather with non-neuronal cell types. Taylor et al. also described this cell type as the one with the least similarity with neuron types (Taylor et al., 2021). Most glial clusters grouped together and resembled neurons, but they also clustered near the hypodermis, consistent with their epithelial hypodermal cell junctions. A smaller group of CEP glial cells exhibited a different and more distinctive transcriptomic signature. Two uterine epithelial cell types, UV1 and UV3 (clusters 146_0 and 68_1) and the spermathecal-uterine valve (sp-ut, cluster 60_0) showed transcriptomes that resembled those of neurons (Figure S1F), despite their morphological differences. UV1 was previously shown to have neuroendocrine functions (Alkema et al., 2005).

### Verification of the cell-type annotation and comparison with public datasets

We tested by microscopy the expression of 19 new marker genes (details below), belonging to a panel of 7 different tissue types (muscles, neurons, seam cells, gland, coelomocytes, glia, hypodermis; Figure 1E, Table S2) by fusing their regulatory regions (promoters, introns and 3’UTRs) to a fluorescent-tag sequence in a plasmid. The large majority of these expression constructs matched their annotation, and marker genes for orphan clusters with putative annotations from the correlation analysis above were expressed at their predicted locations in the animal. We occasionally observed additional unpredicted ectopic locations not reflected in the single-cell data (Table S2), possibly due to a failure of our transgenes to represent endogenous regulation accurately.

We validated the quality of both our expression data and cluster annotations by comparing our data to previously published datasets. Taylor et al. performed scRNA-seq experiments on late larval neurons, and we observed a high level of consistency with our annotated clusters (Taylor et al., 2021). In 88% of the cases, the most correlated cluster annotations in our data and theirs were in agreement (Figure S1G, Table S3). Kaletsky et al. characterized *C. elegans* gene expression in four young-adult tissues (hypodermis, intestine, muscle, neurons) through FACS-sorted bulk RNA-seq (Kaletsky et al., 2018). Our results agreed with theirs for the hypodermis, muscles and neurons. However, we observed less consistent alignment for the intestine (Figure S1H). Intestinal cells have been reported to become polyploid due to endo-reduplication (Hedgecock and White, 1985); thus, some intestinal cells could potentially have been lost during FACS enrichment of diploid cells. Consistent with this, while we observed strong consistency in our intestinal datasets pre- and post-FACS (Suppl. Notes 1), we did observe more and higher-quality intestinal cells in the pre-FACS sample. Thus, we have provided the intestinal markers derived from pre-FACS as well as post-FACS samples in our tabulation (Table S4).

### Novel Cell-Type Specific Gene Expression

Having obtained an unprecedented view of the transcriptionally-defined cell types in the adult worm, we next sought to elucidate the gene expression programs of these cell types. Using differential expression analysis, each cell type was found to express an average of 6,247 genes (with an average expression of transcripts per million, or TPM > 10), of which an average of 669 genes were differentially expressed with log2 fold change (log2FC) > 2 and FDR-corrected q-value < 0.01. To highlight genes that were specific to each cell type and might serve as marker genes for the cell, we identified genes whose expression was enriched more than four-fold in the cluster of interest versus the cluster with the second highest expression (Table S5). In this way, we identified an average of 14.4 marker genes per cell-type cluster. Some of these genes were known cell-specific markers. For example, we recapitulated *dhp-1* as a marker for hypodermis, *ceh-33* for head muscle, *gcy-23* for the AFD neuron and *gcy-14* for the ASEL neuron (Table S5). We also identified many novel markers. For example, the carbohydrate binding protein *clec-166* emerged as a novel marker for M5 neurons, distinguishing them better than does the known marker *vab-15*. We validated M5 neuron-specific expression of *clec-166* in vivo using a fluorescent reporter under the control of the *clec-166* promoter and 3’-UTR (Figure 1E). Conversely, we also computed and added to our dataset the most ubiquitously expressed genes (Table S5).

### Cell-Type Specific Regulation and Transcription Factor Activity

Differential gene expression across cell types is driven by factors that regulate mRNA production and stability. We sought to leverage transcription factor (TF) motif presence in gene promoters to study cell type-specific TF regulation. We first constructed a binary TF-gene potential matrix by identifying 269 known TF motifs from the CIS-BP database in the −500 bp to 100 bp promoter region of each gene. This procedure located an average of 3,378 potential target genes per TF. We quantified TF target activity in each single cell by computing the AUCell statistic for these TF target gene sets in the top 1000 expressed genes.

Noting that many TFs are expressed only in a subset of cell types, we evaluated TF expression and activity across the 211 cell-type clusters. Hierarchical clustering revealed patterns of tissue-specific transcriptional regulation. For example, the homeobox TF *ceh-36* is expressed in a cluster of amphid neurons (AFD, AWCL, ASEL, ASER), and *ceh-36* target-gene activity highlights the same cluster of amphid neurons (Figure 2A, 2B). To define cell-type specific TF activity systematically, we filtered for TFs with cluster-specific expression (log2FC > 2 and FDR q-value < 0.01) and cluster-specific TF motif activity (FDR q-value < 0.01 by Wilcoxon rank sum test). We identified a total of 1,048 tissue-specific associations of the TF expression with its target genes’ activity across the atlas (Table S6). Many TF-cell type relationships are supported by previous publications, such as body-wall muscle specific HLH-1 activity (Figure 2C), seam cell-specific ELT-1 activity, intestine-specific ELT-7 activity, and IL2 neuron-specific DAF-19 activity (Figure S2) (Brabin et al., 2011; Chen et al., 1994; De Stasio et al., 2018; Sommermann et al., 2010).

**Figure 2.**
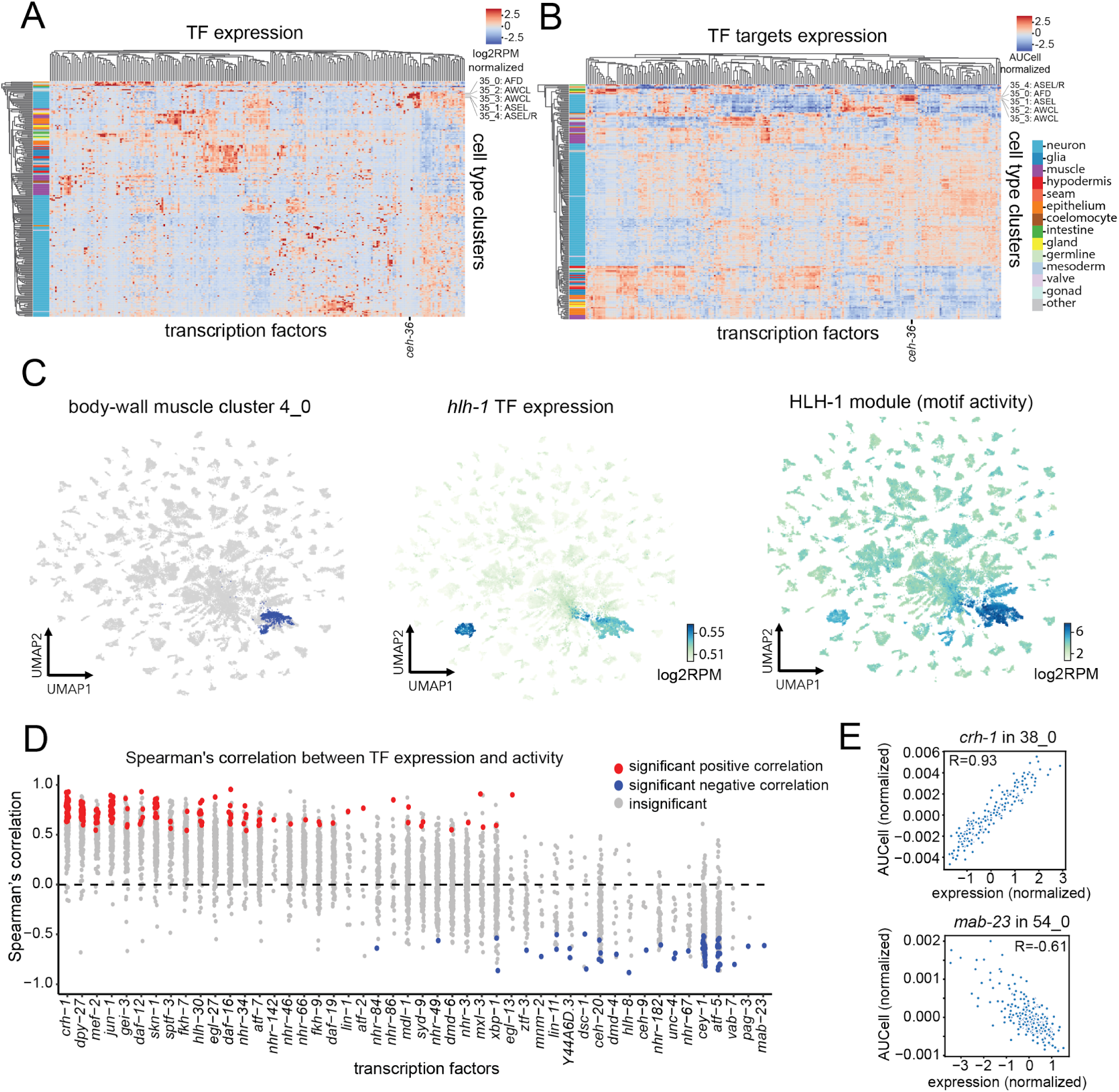
Transcription factor expression and activity across cell types. A) Heatmap showing TF gene expression (log2 reads per million, row-normalized) for 269 TFs (columns) across 211 cell types (rows). Hierarchical clustering was performed on the rows and columns. Cell types are colored by high-level manual annotations. B) Heatmap showing TF activity (AUCell scores of motif targets) for 269 transcription factors across 211 cell types (row-normalized). Cell types are colored by high-level manual annotations. C) HLH-1 activity in the muscle. Left panel: UMAP showing cells belonging to cluster 4_0 (body wall muscle) in blue and other cells in gray. Middle panel: UMAP coloring cells by hlh-1/TFEB TF expression in log2 reads per million. Aside from cluster 4_0, the other cluster with high hlh-1 expression is cluster 23_0, GLR cells. Right panel: UMAP coloring cells by AUCell scores for HLH-1 targets, as defined by motif hits. D) For each of the 48 TFs whose expression and activity significantly correlated in at least one cell type/cluster, we plotted the Spearman correlation of its expression and activity (AUCell score) in each individual cell type. Significant positive correlations by permutation test are in red, significant negative correlations are shown in blue. TFs are ordered by average correlation across all expressed cell types. E) Scatterplot showing correlation between expression and activity (AUCell score) for the crh-1 TF in the intestinal-muscle cluster (38_0) (top); scatterplot showing correlation between expression and activity (AUCell score) for mab-23 in PHC and PVM neuron cluster (54_0) (bottom).

Next, we evaluated the relationship between TF expression and activity in tissues where the TF of interest is expressed. We regressed age as a covariate out of the gene expression matrix to focus on cell-to-cell variation. We then computed the correlation between TF mRNA expression and TF motif activity across the single-cell measurements to focus on examples with strong correlative evidence for a regulatory relationship. We observed a weak positive correlation (Spearman’s r=0.134) averaged across all TF-tissue combinations indicating that on average, TFs play activating roles in gene regulation. TFs with poor expression and activity correlation may be due to RNA abundance serving as a poor proxy for activity, poorly matched database motifs, and/or TF families with many members that bind similar motifs.

We identified 48 TFs whose expression and activity were significantly correlated in at least one cell type (q-value < 0.05 through permutation test). CRH-1, an ortholog of the human activating transcription factor 1 (ATF-1), had significantly correlated expression and activity in 35 cell types. It had an average correlation of 0.55 across all cell types (Figure 2D) and maximum correlation of 0.93 in the intestinal muscle cluster (38_0). Other top TFs consistently showed an activating role (positive correlation between expression and activity) across cell types, for example, DPY-27, MEF-2, JUN-1, GEI-3 and DAF-12 (Figure 2D). We also predicted 19 TFs with repressive activity across cell types. MAB-23 had the most significant negative correlation (R=-0.61) in cluster 54_0 (annotated PHC or PVM neurons). The second most negatively correlated TF was PAG-3 in the muscle cluster 38_0 (Figure 2E). Both TFs contain conserved domains that are linked to transcriptional repression(Inoue and Nishida, 2010; Jafar-Nejad and Bellen, 2004; Lints and Emmons, 2002).

### *C. elegans* Aging Atlas

A prime goal of this study was to define the aging signature of *C. elegans* gene expression. Since all *C. elegans* somatic cells appear to be post-mitotic during adulthood (Sulston and Horvitz, 1977), our dataset is not likely confounded by cell proliferation or replacement. Both shared and cell type-specific gene expression changes have been observed previously during aging, but none of these studies had access to a complete organism (Angelidis et al., 2019; Davie et al., 2018; Enge et al., 2017; Kimmel et al., 2019). Our data enabled the first comprehensive analysis of cell-type aging signatures across an entire animal. We first performed a coarse differential-expression analysis of young cells (day 1, 3 and 5) versus old cells (days 8, 11 and 15) to visualize global trends. Figure 3A displays a heatmap of log2 fold change for the 4,541 genes differentially expressed in at least one cluster, which shows considerable heterogeneity between cell types. On average, each cell type cluster contained 81 genes whose expression changed significantly with age (Figure S3A, Table S7).

**Figure 3.**
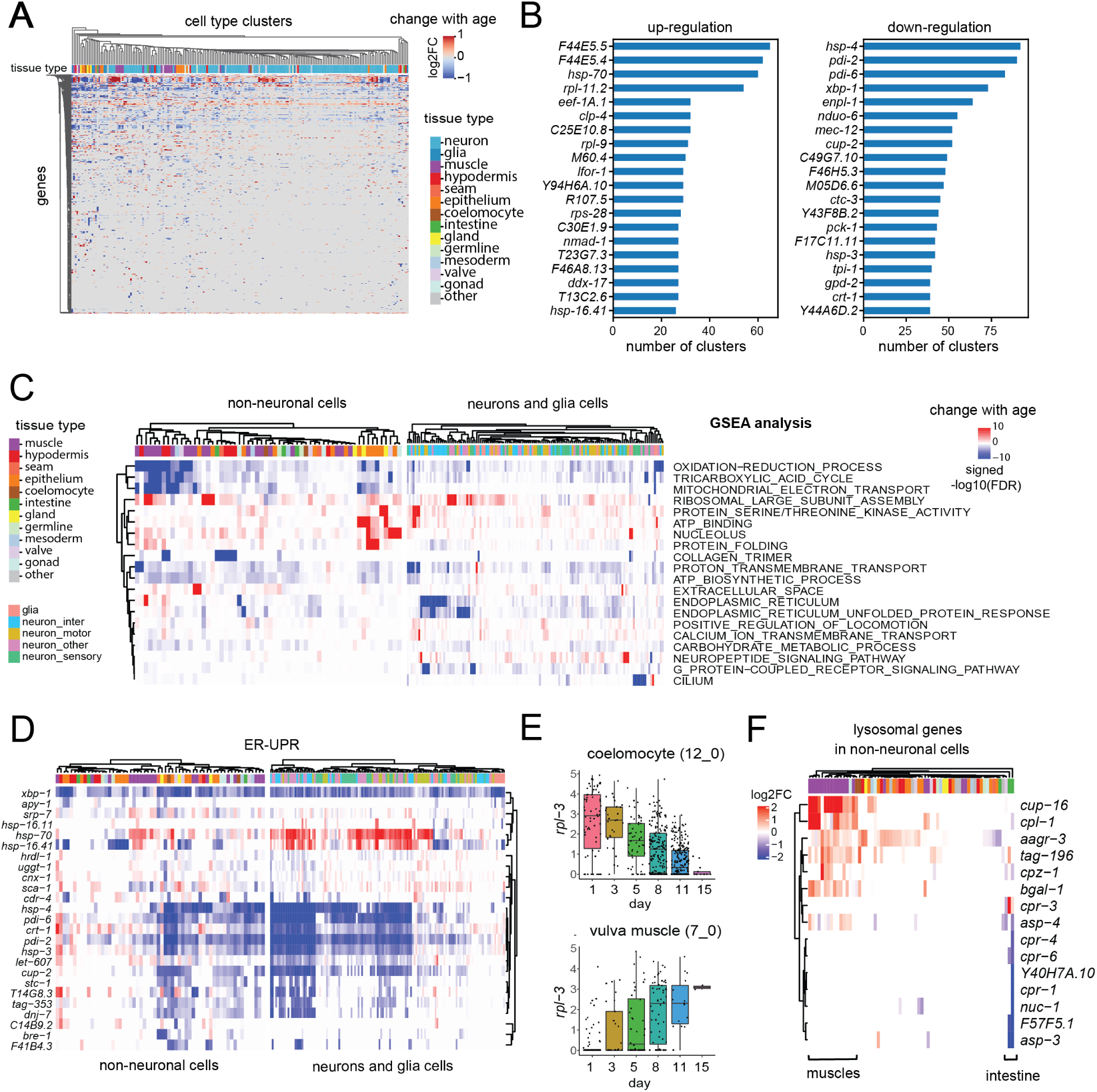
Analysis of shared and tissue-specific aging signatures. A) Heatmap showing log2 fold change (log2FC) in old versus young cells for 4,541 genes differentially expressed in at least one of the 211 clusters. Clusters are annotated by high level cell-type annotations. B) Bar plots show the top 20 differentially-expressed genes ranked by the number of cell types where they are up-regulated (left) and down-regulated (right) C) GSEA results for 20 representative GO terms with significant changes in the most clusters. The rows represent GO terms, and columns represent cell type clusters. The top color bar shows the high-level tissue annotation. Color in the heatmap indicates the signed −log10(FDR q-value) of the GSEA results. Positive indicates that the gene-set abundance is enriched in old cells, whereas negative indicates that the gene set is enriched in young cells. D) Heatmap of differential gene expression log2FC in young versus old for genes in the ^ER^UPR GO term (GO:0030968). The rows represent genes, and the columns represent cell types. E) Boxplot with jitter showing expression of rpl-3 in coelomocytes (cluster 12_0) in log2 (count per 10k) at each time point (top); boxplot with jitter showing expression of rpl-3 with age in vulva muscle (cluster 7_0) in log (count per 10k) at each time point (bottom). F) Heatmap of differential gene expression log2FC in young versus old for lysosomal genes. The rows represent genes, and the columns represent non-neuronal cell types.

We first looked at genes whose expression changes were shared most broadly across cell types. We found that these shared aging genes were more likely to be down-regulated (Figure 3B, S3A), as is the case in mice (Zhang et al., 2021). We ranked the most shared aging genes by the number of clusters in which they were up and down-regulated. Surprisingly, considering our level of detection, we found that regulation of only 10 genes is shared in more than 25% of cell types (Figure 3B). The most frequently up-regulated genes are involved in and may enhance proteostasis. These included the *HSP70* chaperone paralogs F44E5.4 and F44E5.5, orthologs of human *HSPA6*, which were expressed in 30% of clusters (Figure 3B, Table S8). Several additional heat-shock protein genes were frequently up-regulated, including *hsp-70, hsp-110*, *hsp-16.41,* and *hsp-16.2*. Gene set enrichment analysis (GSEA) confirmed that the GO term for heat shock protein binding (q-value = 0.003) was enriched in commonly up-regulated genes. It also revealed that certain cells exhibited GO-term enrichment for ribosome and translation (q-value < 2.2e-16) (Table S9). The most commonly down-regulated aging gene was *hsp-4/BIP,* detected in 44% of clusters*. hsp-4* and the next top three down-regulated genes are involved in the ER-UPR (*hsp-4, cup-2, pdi-6*, *xbp-1*). This suggests either that ER resilience decreases with age or that cells adapt in a way such that these genes are no longer needed. Mitochondrial respiratory chain components (*hsp-3, nduo-6, cox7c, cyc2.1, ctc-3, cox6A*) and ribosomal protein genes (*rps-3, rpl-7A, rps-0, rps-25, rps-9*) were also down-regulated (Table S9). The enrichment of different ribosomal proteins in the up vs. down-regulated sets was curious and is described in greater detail below.

To analyze gene expression patterns at a more functional level, we performed GSEA, this time for each individual cluster based on the log2 fold change in old versus young cells. Out of the 2,166 *C. elegans* GO terms, we found 279 GO terms associated with age in at least one cell-type cluster (q-value < 0.01). We collapsed these to 100 representative GO terms through hierarchical clustering to remove redundancy (see Suppl. Notes 2, Table S10). Consistent with the most shared signatures revealed above, we observed that pathways related to energy metabolism, including mitochondrial respiration, ATP synthesis, glycolytic process and tricarboxylic acid cycle, were the most frequently down-regulated (Figure 3C; Suppl. Notes 2). Overall, we observed that only a small number of GO terms changed globally and consistently across cell types (Figure 3A, S3C).

### Cell-type specific aging regulation

We sought to explore more specific aging signatures that are only observed in single or a subset of cell types. The ER-UPR gene set was down-regulated in several clusters, but in fewer cell types than were the energy metabolism gene sets, and more prominently in neurons compared to non-neuronal tissues (Figure 3D). Conversely, cytosolic chaperones, including *hsp-70, hsp-110, hsp-16.41, hsp-16.2* and *F44E5.4/5*, were up-regulated more strongly past day 3 in a subset of neurons including the amphid neurons (Figure 3D, S3B). Among other gene sets, G-protein-coupled signaling was down-regulated and neuropeptide signaling was up-regulated in the nervous system.

For the large majority of genes, expression changed with age in fewer than 10 clusters (Figure S3C). Among the most noticeable cell-type specific age-related change in our data was the strong movement of ribosomal protein-coding mRNAs with age (Figure 3C, Figure S3B, S3C). Expression of small and large ribosomal subunit genes, as well as translational elongation genes, changed together in either direction depending on the cell type. As an example, the large subunit protein-coding gene *rpl-3* was highly up-regulated in vulva muscles with age but highly down-regulated in coelomocytes (Figure 3E). Previous studies have shown that the overall amount of translation in bulk worm population extracts trends downward with age, correlating with a decrease of bulk ribosomal-protein turnover (Depuydt et al., 2016; Walther et al., 2015) and an increase in ribosomal-protein aggregation (David et al., 2010). Similar mass spectrometry protein turnover analysis revealed heterogeneity among ribosomal protein levels with age (Dhondt et al., 2017) that could be linked to the cell-type heterogeneity that we observe in our expression dataset.

A closer examination of the cell-type specific GSEA results revealed additional tissue-specific pathway changes (Suppl. Notes 2). Cuticular collagen matrix genes (GO:0005581) were significantly down-regulated specifically in the hypodermis, seam cells, phasmid sheath glia, and one intestinal cluster, whereas another subset of extracellular matrix genes coding for basal membrane constituents was down-regulated in muscle cell types (Suppl. Notes 2). The intestinal cells down-regulate several lysosomal protease genes with age (*cpr-1/4/6*, *asp-3/4*, *cpl-1*, *cpz-1*), whereas many muscles up-regulate lysosomal genes, including proteases and enzymes predicted to be involved in polysaccharide and lipid catabolism (*asp-4*, *cpl-1*, *cpz-1, spp-10*, *bgal-1*, *cpl-1*) (Figure 3F). Notably, increased activity of GLB1, the homolog of the beta-galactosidase *bgal-1* in worms, is a conserved hallmark of cell senescence in mammals (Dimri et al., 1995). The proteasome subunit genes represented another example of tissue-specific aging dynamics; they were stable in most cell types but were up-regulated in several muscle clusters and down-regulated in the somatic gonad and some uterine epithelial cells (Suppl. Notes 2).

To extend and complement our GO-term based analysis of aging signatures described above, we also looked at the activity of *C. elegans* gene sets known to be co-regulated across a wide range of perturbations. To this end, we used a collection of 209 gene expression modules described previously (Cary et al., 2020) and derived using Deep EXtraction Independent Component Analysis (DEXICA) of a large, diverse collection of *C. elegans* microarray experiments. We asked which DEXICA modules changed the most with age (Figure S3D). Some of the most commonly age-dependent modules were related to ribosomal biogenesis, collagen synthesis, the ER-UPR and mitochondria (Figure S3D), confirming some of the GO-term results described above. Interestingly, this analysis also highlighted DNA damage response and DNA repair, which were not identified by GO-term analysis (Figure S3D). This gene set was significantly and mostly monotonically up-regulated with age in 35 clusters, as exemplified by motor neuron and seam cell clusters (Figure S3E).

### Dynamic Gene Expression Changes with Age

To characterize dynamic gene expression changes with better temporal resolution, we studied trajectories for the 4,232 genes significantly differentially expressed in the young versus old comparison in at least one cluster (|logFC| > 1 and q-value < 0.01). We computed an average trajectory for each gene across cell types and grouped the genes into six age modules through hierarchical clustering as in (Schaum et al., 2020). We then associated each gene module with GO terms by GO enrichment analysis (Figure 4A, S4A, Table S11).

**Figure 4.**
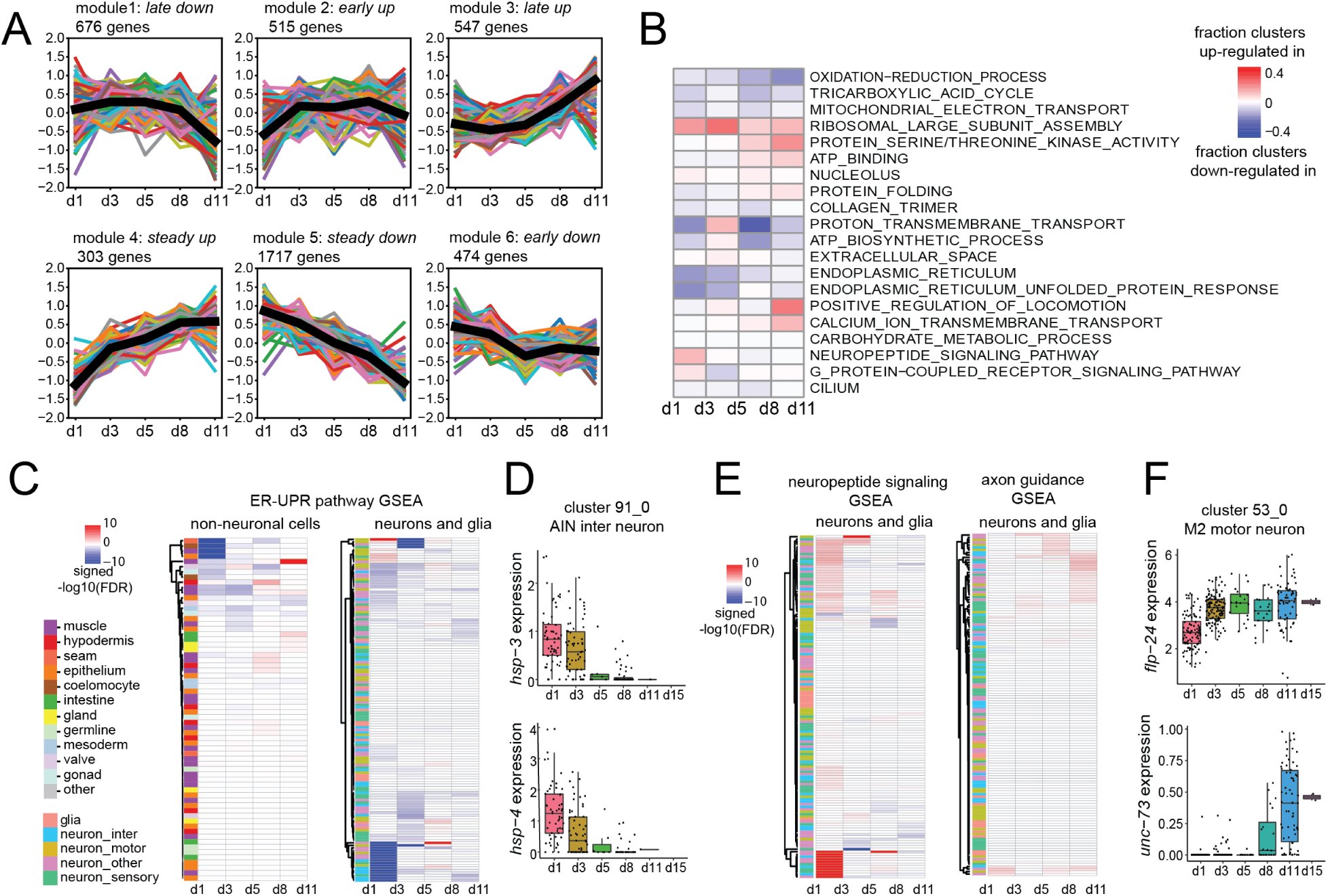
Dynamic gene expression analysis with age reveals six major temporal trajectories. *A)* In each panel, a colored line represents the average trajectory of genes in that module for a particular cell type and the black line represents the averaged trajectory over all cell types. Gene modules are defined as genes showing a similar aging trajectory (Methods). *B)* Heatmap showing the fraction of clusters in which the corresponding GO term (rows) is significantly up-regulated (red), or significantly down-regulated (blue) with age between each consecutive time point pair (columns). *C)* Heatmap showing the GSEA result for the ER-UPR (GO:0030968), measured by signed −log10(FDR) between each pair of consecutive time points for each cell type (left, non-neuronal cells; right, neurons and glia). *D)* Boxplot with jitter showing expression of the ER chaperone genes hsp-3 (top) and hsp-4 (bottom) in the AIN interneuron (cluster 91_0) in log (count per 10k) at each time point (top). *E)* Left: heatmap showing the GSEA result for neuropeptide signaling (GO:0007218) measured by signed −log10(FDR) between each pair of consecutive time points for neuron and glial cell types; Right: heatmap showing the GSEA result for axon guidance (GO:0008045) measured by signed −log10(FDR) between each pair of consecutive time points for neuron and glial cell types. See Figure S4B for results for non-neuronal cells. *F)* Boxplot with jitter showing expression of flp-24 (top) and unc-73 (bottom) in the M2 motor neuron (cluster 53_0) in log (count per 10k) at each time point (top).

The six age modules showed diverse aging trajectories. Modules 3 and 4 contained genes up-regulated with age. Module 4 contained immune response genes that showed an early adulthood up-regulation that stabilized later in life. In contrast, module 3 genes (enriched for axon guidance and neuronal development) were up-regulated only in late life. Modules 1, 5, and 6 contained genes that were down-regulated with age, with different trajectory shapes. Genes in module 5 (mitochondrion, oxidation-reduction) steadily declined with age, while genes in module 6 (ER-UPR) and 1 (translation) declined early and late, respectively. Module 2 was also enriched for translation genes. While these genes exhibited an early increase with age, the trend reversed in late age (Table S11).

Next, we explored the time-series data from a pathway-centric view. We performed differential expression analysis followed by GSEA for cells in consecutive time points in each cell type cluster. Figure 4B shows the result for the same pathways as Figure 3C with finer time-scale resolution. Consistent with gene trajectory analysis, we observed that oxidation-reduction processes underwent a consistent decline throughout the aging process, whereas the neuropeptide signaling pathway increased sharply during days 1-3.

For specific pathways of interest, we then explored the pathway-change dynamics in each cell-type cluster. For example, we found that the ER-UPR pathway was significantly down-regulated only during days 1-5 in specific subsets of neuronal and non-neuronal clusters including the epithelium, coelomocytes and muscles (Figure 4C). As an example, we examined *hsp-3* and *hsp-4* mRNA levels in AIN interneurons (cluster 91_0) and confirmed that they indeed experience the most dramatic decrease early in aging (Figure 4D). Pathway dynamic change analysis also revealed early increase of neuropeptide pathways during aging and late increase of several gene sets involved in neuronal growth like axon guidance and neuron projection morphogenesis pathways (Figure 4E). We examined the expression of specific genes in M2 motor neurons (cluster 53_0). Neuropeptide *flp-24* mRNA levels increased dramatically from days 1-3, while mRNA of *unc-73*, a key gene for axon guidance (Steven et al., 1998), only started to increase at day 8 (Figure 4F). This observation is consistent with previous reports describing ectopic neurite branching in old neurons (Pan et al., 2011; Tank et al., 2011; Toth et al., 2012).

In conclusion, our aging atlas revealed that the decline of carbon and mitochondrial metabolism is the most consistent aging signature, occurring across many cell types and continuously during the entire course of aging. Other functions including ER-UPR, ribosome, lysosome, proteasome and collagen trimers shifted significantly with age in a cell-type-specific or tissue-specific fashion. Moreover, the temporal resolution of our dataset enabled us to identify the distinct aging trajectories of various pathways.

### Global Transcriptional Characterization Reveals Cell-Type Specific Aging Patterns

Gene expression drift is a common correlate of aging (Kimmel et al., 2020; Rangaraju et al., 2015; Tarkhov et al., 2019). We sought a full transcriptome quantification for how extensively gene expression changes with age in each cell type. We first quantified the average magnitude of transcriptome change between consecutive time points by maximum mean discrepancy (MMD) (Kimmel et al., 2020) and Methods). MMD is a statistical test measuring the distance between two distributions (Gretton et al., 2012); here, the cell embeddings in different time points. We assume that the degree to which MMD deviates with time is a proxy for how much each cell type is altered by aging, and we refer to it as the aging magnitude.

For cell-type clusters with enough cells to evaluate MMD changes between consecutive time points, we observed significant changes in at least two consecutive time points at a threshold of FDR q-value < 0.01 for 158/165 cell types. We verified that the MMD metric was independent of the number of cells in the cluster (Figure 5A, S5A). More broadly, cell types with similar ontological relationships and baseline transcriptional profiles had more similar aging magnitudes, suggesting the hypothesis that cellular function influences aging rate (Figure S5B). For 7 cell types, changes during aging were insignificant by the permutation test (Table S12). Interestingly, the magnitude of this change for the other 158 clusters exhibited substantial heterogeneity both between different cell types and within a single tissue type (Figure 5A). An amphid neuron cluster (63_0) changed the most during aging, whereas cluster 3_0 (related to germline) changed the least (Figure S5C). In general, neuron clusters exhibited a significantly greater aging magnitude than did non-neuronal clusters (p < 1.64e-06, MMD Wilcoxon rank sum test, Figure 5B).

**Figure 5.**
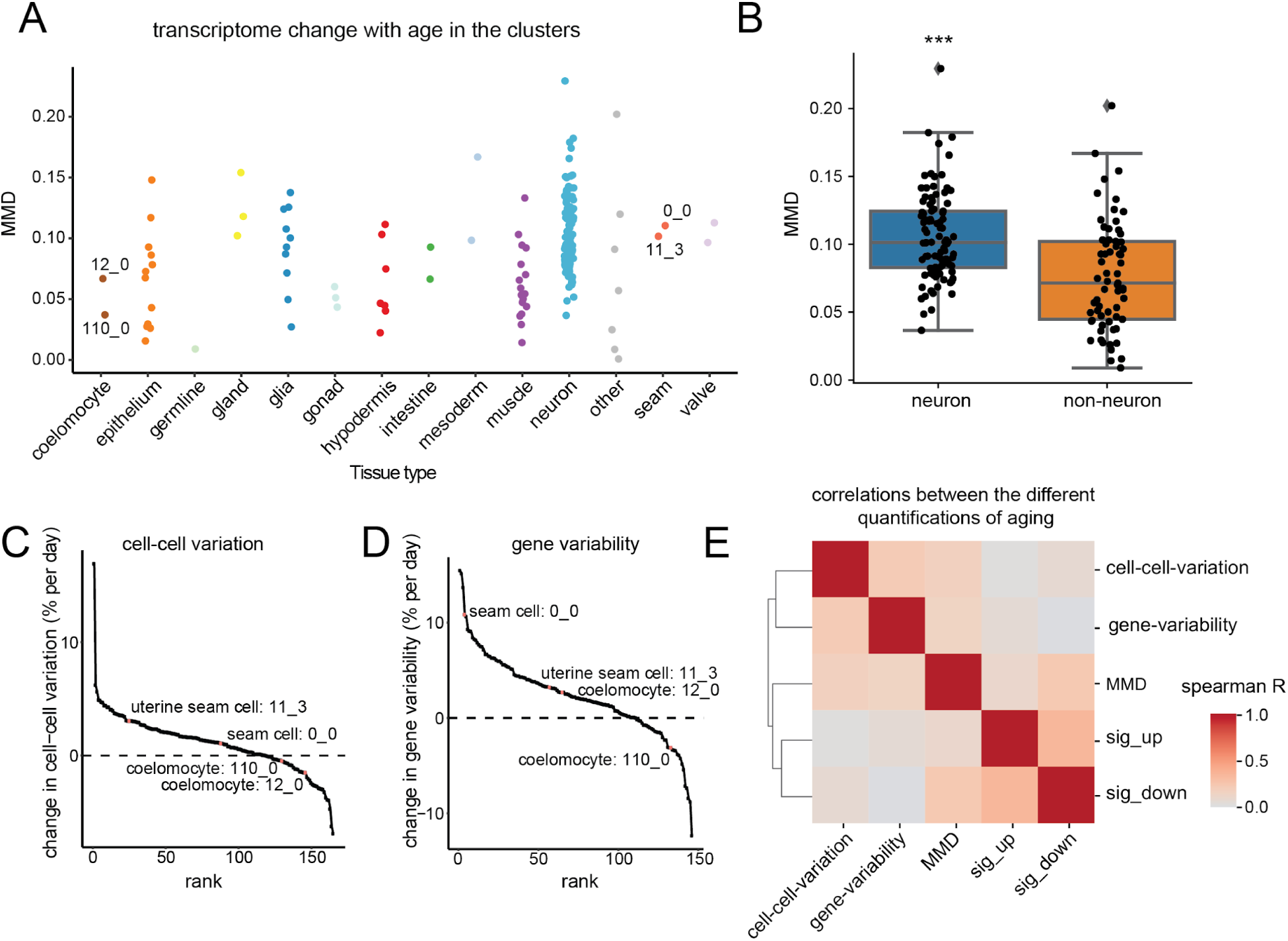
Global aging characterization reveals difference in magnitude, cell-cell variation, and gene variability during aging across cell types. *A)* Transcriptome changes during aging estimated by MMD (maximum mean discrepancy) for all cell type clusters. For each cell type cluster, we first computed MMD between each consecutive pair of time points (d3-vs-d1, d5-vs-d3, d8-vs-d5, d11-vs-d8 and d15-vs-d11), and reported the mean MMD across all time points. Coelomocyte clusters and seam cell clusters are labeled. Clusters 12_0, 110_0, 11_3 refer to the cell types analyzed by microscopy in Fig.6. *B)* Boxplot comparing MMD estimates in neurons versus non-neuronal cell types. MMD statistics are significantly larger for neurons compared to non-neuronal cells (*** p < 1.64e-06, Wilcoxon rank sum test). *C)* Cell-type clusters ranked by rate of change in cell-cell variation (% per day) during aging. Coelomocyte clusters and seam cell clusters are labeled. *D)* Cell-type clusters ranked by their percentage change in variance of Pearson residual (% per day), which quantifies gene expression variance. Coelomocyte clusters and seam cell clusters are labeled. *E)* Heatmap of Spearman’s correlations between the different quantifications of aging in the transcriptional space across cell type clusters. MMD, cell-cell variation, expression variability, sig_up, average increase in gene expression for 34 commonly up-regulated genes; sig_down, average decrease in gene expression for 114 commonly down-regulated genes.

Complementary to global transcriptional changes, prior studies have observed increased variance in mRNA abundance with age (Martinez-Jimenez et al., 2017). We considered two distinct strategies to quantify variance in our data and compare across age. First, for each cell type cluster, we focused on the range of expression profiles observed by computing the average Euclidean distance from each cell to the population centroid at each time point (cell-cell variation). We observed increased cell-cell variation in 116 clusters, but surprisingly, we also observed reduced cell-cell variation in 49 clusters (Figure 5C). Thus, cell-to-cell variance increases in the majority of cell types but can narrow with age in many cell types. Similar observations were made comparing the cell types from several organs of young and old mice (Kimmel et al., 2019).

Second, we focused on the variability of each individual gene, which could reflect large transcriptional bursts rather than steady, consistent transcription. We quantified the biological variability in mRNA abundance (gene variability) within a homogenous cell population (same age, same cell type) by fitting a negative binomial regression model to each gene and computing the variance of the analytical Pearson residual (Hafemeister and Satija, 2019; Lause et al., 2020). The Pearson residual is intuitively a measurement of goodness-of-fit with respect to a null distribution, assuming all cells share the same transcriptional profile (Lause et al., 2020). To focus on variance at the cell-type level, we summed every gene’s statistic for each cell type cluster. Out of the 146 cell-type clusters for which we have sufficient data from each time point, we observed 110 cell-type clusters with increased expression variance during aging and 36 with reduced variance (Figure 5D). Thus, we observe that most cell types show increased variability with age, consistent with previous work (Bahar et al., 2006). Interestingly, we observed that neurons not only show a greater change in terms of MMD measurement, but they also tend to show a greater change in cell-cell variation and gene-variability with age (Figure S5D).

Finally, we explored the relation across these various approaches to quantifying global aging magnitude across the different cell types (Table S13). We included MMD transcriptome change, cell-cell variation, and gene variability. Given the intriguing set of metabolic genes and chaperones observed to change with age across many cell types above, we also quantified this core aging signature in each cell type cluster by looking at the average log fold changes of a set of shared differentially expressed genes between young and old cells. We named Sig_up (or Sig_down) the genes observed in more than 50 clusters that have an average log2FC > 0.5 (or log2FC < −0.5). Interestingly we observed weak positive correlations among the distinct statistics (Figure 5E, S5E), suggesting that aging results in a global transcriptional change that can be observed across multiple axes with concordance.

We hypothesized that cell types that trended toward greater or lesser MMD values across multiple statistics could be more reliably highlighted as faster- or slower-aging cells. Previous studies in worms have reported that late-life muscle mitochondrial fragmentation is associated with health decline and predicts increased mortality (Hahm et al., 2015; Roux et al., 2016). Thus, we asked whether the extent of mitochondrial fragmentation *in vivo* might correlate with the magnitude of age-specific gene expression changes. We selected uterine seam cells (clusters 11_3) and two coelomocyte clusters (clusters 12_0 and 110_0) for this experiment. In the transcriptional space, uterine seam cells appear to age faster than coelomocytes, supported by greater MMD (0.10 vs. 0.07 and 0.04), cell-cell variation (3.04 vs. −1.54 and −0.47), and gene variability (3.23 vs. 2.66 and −3.12). Using fluorescent microscopy, we scored age-related change in mitochondria morphology concomitantly in the coelomocytes and the uterine seam of many individuals (Figure 6A). We observed that, on average during aging, the uterine seam cells have more fragmented mitochondria than the coelomocytes (Figure 6A, 6B, S6B), which in turn retain a more youthful tubular mitochondrial network compared to the uterine seam cell (Figure 6C, S6C). In addition, we compared aging mitochondria in two other cell types, body wall muscles (cluster 4_0 and 4_1) and excretory glands (cluster 89_0) (Figure S6D-F). Our observations confirmed again a concordance with the changes in magnitudes of gene expression of these two cell types. Together, these *in vivo* studies support the hypothesis that the larger changes in overall gene expression between early and late time points reflects a faster cellular aging rate.

**Figure 6.**
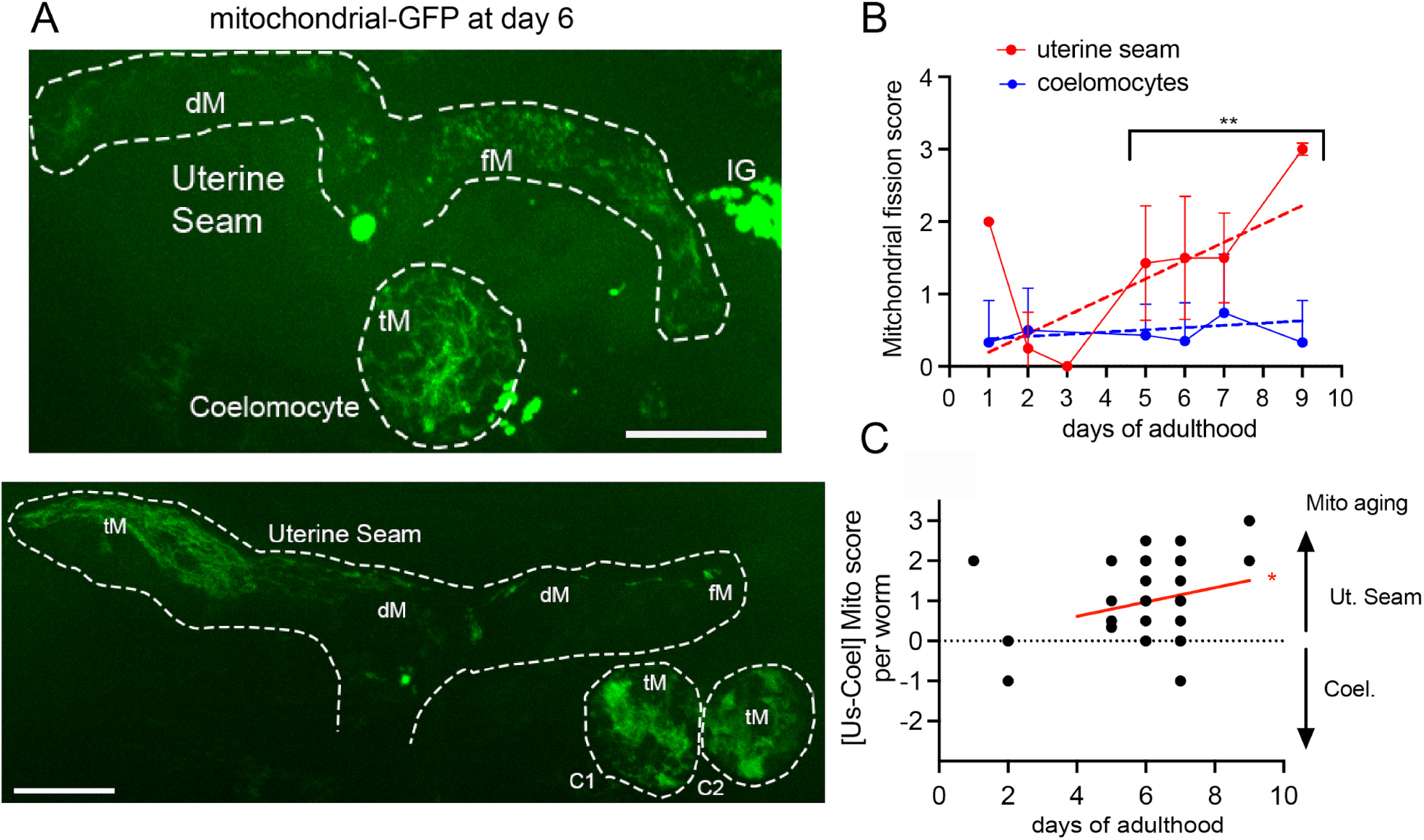
Quantification of mitochondrial morphology changes with age in two cell types corroborate the rates of change measured by gene expression. *A)* Two representative images showing mitochondrial morphology at mid-age in coelomocytes (cluster 12_0) and uterine seam cells (cluster 11_3), which exhibit a small and large magnitude of aging, respectively, according to expression data. GFP expression was driven by tissue-specific promoters and the protein was targeted to the mitochondrial matrix. Additional images of young and old cells are shown in figure S6A. dM: depleted mitochondria. fM: fragmented mitochondria. tM: tubular mitochondria. IG: autofluorescent intestinal granules. C1,C2: Coelomocytes 1 and 2. Scale bar: 10𝝻m *B)* Mitochondrial fragmentation score during aging in coelomocytes and uterine seam cells. ** p=0.01. Dashed lines: linear regression. *C)* Comparison of coelomocytes and uterine seam mitochondrial fragmentation score in the same animal during aging. * linear regression different from zero p=0.04.

### Transcription Factor Regulation during Aging

Transcription factors play a crucial role in the regulation of longevity across many organisms (Kenyon, 2010). Assessing TF activity with single-cell resolution across age can uncover new relevant regulators that bulk tissue analysis may have missed, since most TFs act in a limited number of cells (Figure 2A). We summarized TF expression change across age for each cluster by its log2FC and target gene change by a t-statistic comparison of target gene AUCell scores in young versus old cells. We observed that for 22 putative activators (Figure 2D), TF targets changed expression in the same direction as the TF itself during aging (mean correlation = 0.35, Figure S7A). For 17 putative repressors (Figure 2D), TF targets changed in the opposite direction as the TF itself during aging (mean correlation = −0.26, Figure S7A).

Figure 7A shows the top 20 TFs with the most positive average log2FC across cell types (see Figure S7A, S7B for changes in AUCell scores). Remarkably, of the eight top most globally age-regulated TFs, seven are well-known positive regulators of *C. elegans* longevity listed in the GenAge database (of the Human Aging Genomic Resources): *skn-1, daf-16, hlh-30, fkh-7, fkh-9, daf-12, dpy-27* (Kaletsky et al., 2016; Tacutu et al., 2018). Two other TFs that are consistently up-regulated but in fewer cell types, *nhr-23, nhr-25* are longevity regulators as well (Tacutu et al., 2018). We observed that *skn-1/Nrf2*’s age-dependent expression change showed interesting cell type specificity; whereas *skn-1/Nrf2* expression increased in most cell types, it decreased in glial cells. Likewise, for some glial cell type clusters, the *skn-1/Nrf2* expression decrease was accompanied by a decrease in its predicted targets as well (Figure 7C). *gei-3* is a broadly expressed TF that has not previously been associated with aging. We found universal up-regulation of *gei-3* and its predicted targets (Figure 7D), suggesting that it could be an interesting new longevity regulator.

**Figure 7.**
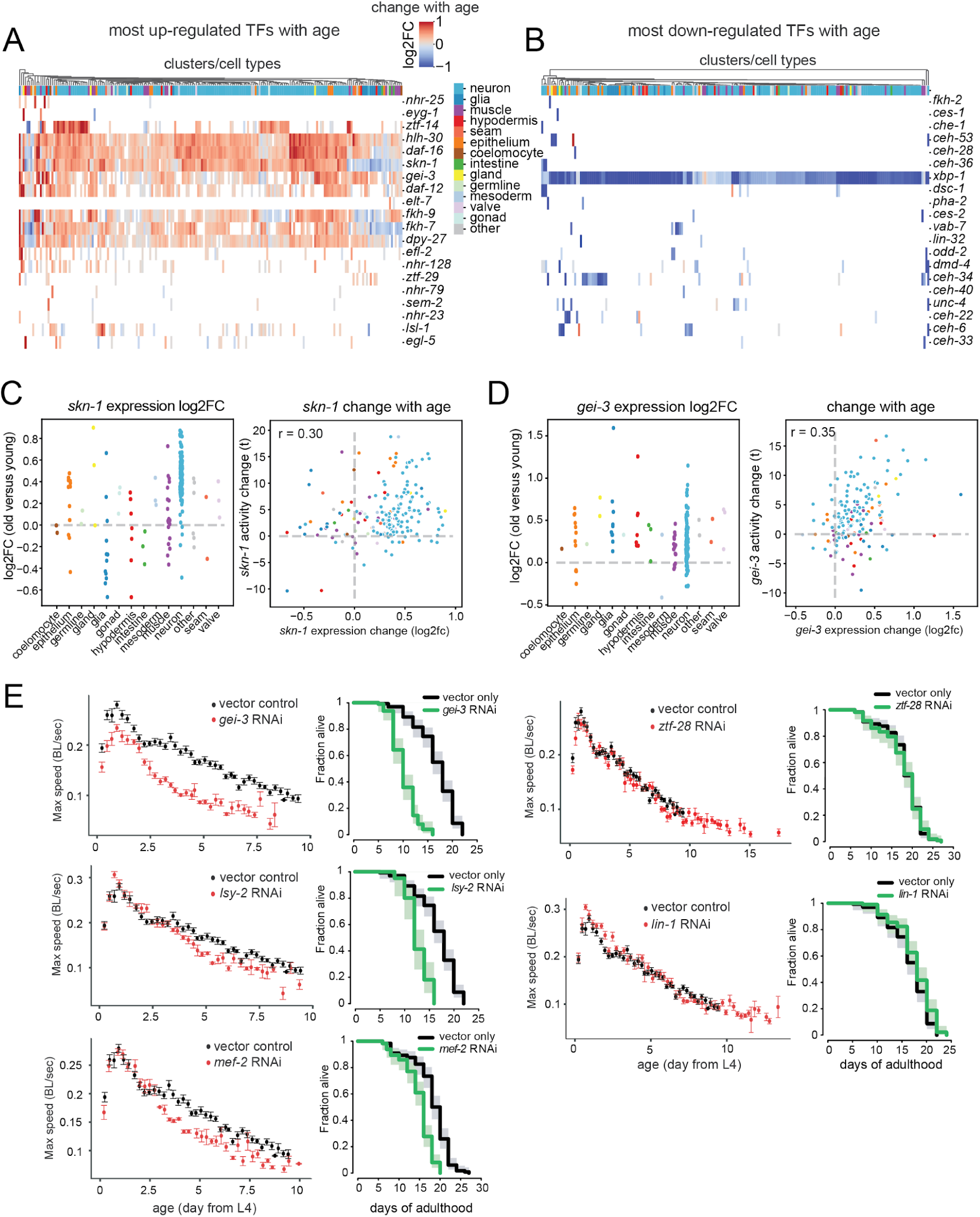
Changes in transcription factor expression and activity during aging. *A)* Heatmap of TFs expression log2FC during aging for top 20 TFs with the most positive average log2FC across cell types in which they are expressed. A blank indicates that the TF is not expressed in that cell type. *B)* Heatmap of TFs expression log2FC during aging for top 20 TFs with the most negative average log2FC across cell types that they are expressed in. A blank indicates that the TF is not expressed in that cell type. *C)* Left panel: expression log2FC of skn-1 in each cell type cluster. Right panel: scatterplot of the expression log2FC and changes in AUCell scores for skn-1. *D)* Left panel: expression log2FC of gei-3 in each cell type cluster. Right panel: scatterplot of the expression log2FC and changes in AUCell scores for gei-3. *E)* Left panels: decline in maximum speed (healthspan) after RNAi treatment targeting transcription factors up-regulated with age in many tissues. (red curves). Right panels: Survival after RNAi treatment targeting the same genes as in A. (green curves).

Figure 7B shows the top 20 TFs whose expression decreases the most during aging. In contrast to the up-regulated TFs, the majority of down-regulated TFs were cell type specific. Only *xbp-1* and *unc-4* are known lifespan regulators (Tacutu et al., 2018). *xbp-1* mRNA abundance decreased universally across cell types. *xbp-1* is a key regulator of the ER-UPR (Imanikia et al., 2019), which we also observed to have decreased expression as a gene set with age in many cell types (Figure 3F, 4C). In summary, TF expression changes with age often, but not always, correlate with the activity of their targets. Moreover, we observed that TFs universally up-regulated with age across many tissues are enriched with TFs whose normal function is known to extend lifespan.

### microRNA regulation during aging

In addition to TFs, microRNAs, which can trigger mRNA degradation(Bartel, 2004), have been explored as longevity regulators in *C. elegans* (Pincus et al., 2011). We examined the enrichment of miRNA targets for genes differentially expressed during aging for each cell type cluster for the 60 miRNA families in the targetScan database (Methods). The miRNAs had an average of 157 predicted targets. We found 16 miRNA families whose targets are significantly enriched (q-value < 0.01) for differentially expressed genes with age in at least one cell type (Figure S7C, Table S14), four of which were reported to change in expression during aging in a previous bulk study (Lencastre et al., 2010). Our result is consistent with a previous finding that the majority of miRNA targets are up-regulated with age, suggesting that their expression is de-repressed following the decline of miRNA levels(Ibanez-Ventoso and Driscoll, 2009). Among them, miR-34 and miR-238 are known to be involved in regulation of *C. elegans* lifespan (Lencastre et al., 2010; Yang et al., 2013). This analysis showed that a subset of gene expression changes observed during aging is likely due to changes of microRNA levels, and that the majority of these are regulated in a cell-type specific way.

### Screen for Novel Longevity Transcription Factors

Based on the observation that TFs whose expression changes in many tissues during aging are enriched for longevity regulators, we investigated 55 TFs whose expression (or target activity) changed with age but were not linked previously to aging (Figure 7A; Table S15). We used RNA interference (RNAi) to turn down their expression starting from early adulthood and systematically recorded animal movement rates and survival over time. To do this, we adapted a real-time computer vision “Worm Tracker” (Swierczek et al., 2011) to an automated, high throughput format (Kerr et al. 2022).

We found that RNAi of three such TF genes, *gei-3*, *lsy-2*, and *mef-2* accelerated the rate of movement decline and mortality (Figure 7E). *gei-3* RNAi produced the largest effect, similar to that of RNAi inhibition of a positive control longevity regulator, the heat shock TF *hsf-1* (Figure 7E, S7D). *lsy-2* and *mef-2* RNAi produced similar but less dramatic effects both on lifespan and movement (Figure 7E). Importantly, RNAi initiated at hatching did not affect developmental speed nor survival of larvae (Figure S7E). The survival phenotype only appeared after the worms reached adulthood, suggesting these TFs are genuine longevity regulators. In addition, we identified two TFs, *lin-1* and *ztf-28*, whose RNAi extended healthspan. We recorded increased movement speed compared to control consistently towards the end of life, a period characterized by a lethargic state (Zhang et al., 2016). Upon these RNAi treatments, a subset of animals could pass the movement detection threshold for up to 30% longer than the control. Surprisingly, neither *lin-1* nor *ztf-28* RNAi affected lifespan (Figure 7E), suggesting a decoupling of mortality and health decline. We observed a similar phenotype for the knockdown of another positive control, the translation initiator *eif-1,* (Figure S7D) which was previously reported to increase lifespan in other conditions (Curran and Ruvkun, 2007). While little is known about *ztf-28*, *lin-1* is a well-studied gene involved in vulval cell fate determination during development (Beitel et al., 1995). Our analysis shows that increased *lin-1* expression with age is more prominent in the muscles, possibly leading to a detrimental effect on movement. All five of these TF hits have homologs in the human genome.

## Discussion

### Resource

In this study, we present a single-cell atlas of the adult nematode *C. elegans*. To date, the Tabula Muris Senis consortium has offered the most comprehensive view into the gene expression changes at the cellular level in many cell types and organs (Tabula Muris Consortium et al., 2018; Zhang et al., 2021). Due to the small size and cell number of *C. elegans*, using this organism, we could extract virtually every cell type from a large number of animals, in parallel in the same laboratory, reducing the variation between the tissues and time points due to non-biological factors. The dataset provides an original and near-complete resource for cell-type specific expression in *C. elegans* adulthood and a history of gene expression changes that accompany aging. The quality of our dataset was assessed and validated both by comparison to imaging experiments and to previous scRNA-seq of larval stages.

Regulatory sequences for cell-type specific marker genes are an invaluable tool to genetically target and perturb a given cell type. Their identification has historically been empirical and often fortuitous (Dupuy, 2004; Hope, 1991). This scRNA-seq dataset provides a new collection of adulthood marker genes for every cell type, including many cell types previously lacking such specific promoters. To enable additional analyses, we have made the entire dataset accessible through an online interface. Users can query the cell-type specific expression of single genes, GO terms or custom gene sets. They can also explore gene expression changes during aging. Finally, we have made our analysis of TFs and their target gene sets across cell types and age available through the interface.

### Aging signatures and TFs

Our analysis brings to light a high level of heterogeneity of coherent gene expression changes with age between cell types. The list of genes whose expression changes broadly across many clusters/cell types during aging is surprisingly short (Figure 3A, S3C). For instance, we observe that only 12 genes change with age in more than 50 cell types/clusters (Figure 3B). To some extent, this may be explained as different cell types express different cell identity genes whose age-dependent changes could be cell type specific. Consistent with this, the more global gene changes largely involve broadly expressed metabolic and other “housekeeping” genes. The most common aging signature that we observe is a down-regulation of cytoplasmic and mitochondrial carbon metabolism genes, as well as mitochondrial respiration genes (Figure 3B, 3C). The decline of mitochondrial respiration is one of the most conserved aging phenotypes observed across distant phylogenies (López-Otín et al., 2013), including *C. elegans* (Brys et al., 2010).

That said, we observed considerable, non-random variation in expression of genes that do not define cell identity but rather processes involved in molecular repair and resilience. For example, we observed frequent down-regulation of ER genes, including ER stress response, occurring more widely in the neurons (Figure 3D). This discovery complements the discovery by Kimmel et al. in mice that the most commonly down-regulated GO terms in aging kidney, lung, and spleen comprise proteins targeted to the ER (Kimmel et al., 2019). These observations across distant species suggest that down-regulation of ER functions could be a recurrent aging mechanism, perhaps a causal one. Our analysis found that the loss of ER stress response gene expression is an event limited to early aging. It could explain the early decline of *C. elegans’* ability to respond to ER stress (Ben-Zvi et al., 2009). Also among our common up-regulated aging signatures were cytosolic chaperones including small heat shock proteins and the HSP-70 family (Figure 3B, S3B), as previously seen in bulk analysis (Lund et al., 2002). The gradual demise of proteostasis with age is a common hallmark of aging (López-Otín et al., 2013) and has been reported by several methods in worms (Ben-Zvi et al., 2009; David et al., 2010), suggesting that these chaperones are up-regulated as a compensatory process that may help to restore homeostasis. How each tissue is differently affected has not been reported previously. Our scRNA-seq indicated a particularly strong up-regulation of chaperone gene expression in the neurons (Figure 3D, S3B), suggesting this tissue is subjected to higher protein stress with age. Likewise, we find a strong DNA-repair signature in specific cell types including neurons, valves, and epithelium cells (Figure S3D).

The *xbp-1* transcription factor controls one of the branches of the ER-UPR stress response and is among the most broadly down-regulated genes with age (Table S8). Frakes et al. found that restoring *xbp-1* activity specifically in some glia cells can trigger a signal that activates the ER-UPR stress response in the peripheral tissues, like the intestine, and in turn increases longevity (Frakes et al., 2020). Interestingly, we found that glia clusters experienced the strongest drop of *xbp-1* mRNA with age, suggesting that this loss could drive aging (Figure 3D, S3B).

When considering aging signatures in each cell type at the level of functional GO terms, we found that the great majority of gene expression changes are cell-type specific or only shared in a limited subset of tissues (Figure 3A). Signatures including the ribosomal genes or the protein folding stress response are present in many cell types but not universal. Others, for example, genes encoding proteins associated with the extracellular matrix, lysosome, innate immunity, proteasome, DNA repair, or neuronal growth, are more specific and linked to the intrinsic functional specificities of the cell types (Figure 3). Ribosomal mRNA trajectories are among the most differentially regulated signatures of our dataset, exhibiting a strong down-regulation in some cells but an up-regulation in others (Figure 3E). A similar effect is also emerging from the analysis of scRNA-seq data of aging mice (Zhang et al., 2021). Interestingly, though this effect is different, even reversed, between different cell types, it is consistent within the cell population of a cell type cluster, suggesting that the cause of this change in mRNA abundance is not haphazard but rather part of a cell-type program. Different cells could change expression of different gene sets depending on their specific functions (gut barrier, secretory, metabolic, mechanosensory, innate-immunity and so forth). These coordinated gene changes may influence the rate of aging; for example by increasing proteostasis or repair or even by reducing ribosomal protein levels, as was shown experimentally in yeast and in worms (Hansen et al., 2007; Kaeberlein et al., 2005).

One of our most surprising discoveries was that canonical life-extending TFs together comprise the most broadly up-regulated TFs with age. These included *daf-16/FOXO3*, *hlh-30/TFEB*, and *skn-1/NRF2* (Lapierre et al., 2013; Martínez Corrales and Alic, 2020), three conserved stress-response TFs, as well as *daf-12*, *dpy-27, fkh-9 and fkh-7,* which have also been found to promote longevity in worms (Hsin and Kenyon, 1999; Kaletsky et al., 2016; Mansfeld et al., 2015; Tacutu et al., 2012). These pro-longevity genes are all known stress-response TFs, suggesting that their up-regulation occurs in response to cellular stress, potentially alleviating the cellular burden of aging and preventing it from causing damage even sooner.

By testing other TFs whose age-dependent expression changes similarly in multiple cell types, we found five new TFs influencing lifespan and/or healthspan. *gei-3,* homologous to CIC (Capicua) in mammals, is altered across most cell types during aging (Figure 7D) and has the strongest effect on lifespan and healthspan (Figure 7E). CIC was shown to function as a tumor suppressor in two different cancer models (Bunda et al., 2019; Lee et al., 2020). Our other hit, LIN-1 belongs to the eukaryotic ETS-domain TF family. Recently, expression data from long-lived people observed lower levels of ETS1 TF (Xiao et al., 2022). Our result validates the use of scRNA-seq to find new TFs that influence longevity and suggests an examination of TFs whose expression changes globally with age in other organisms.

### Magnitude of Aging

We characterized the global aging pattern of the transcriptome in every tissue by looking at the maximum mean discrepancy (MMD), cell-to-cell variation and gene-variability. Such methods were used previously (Bahar et al., 2006; Kimmel et al., 2020; Martinez-Jimenez et al., 2017), but have not been compared carefully to each other or validated experimentally. Our analysis revealed increased MMD for 96% of the cell type clusters, increased cell-cell variation in 70% and increased gene-variability in 75% of the cell type clusters. This suggests that the majority of cells change both their baseline transcriptome profile and cell-to-cell variance during aging. We observed a modest correlation between these independent measurements, suggesting that some are coordinated (Figure 5E). Age-dependent changes in mitochondrial morphology *in vivo* were used as a proxy to measure aging rate independently. In two separate experiments, we compared the mitochondrial morphology change of two cell types showing different transcriptional aging patterns (Figure 6, S6). In both cases, we found that cell types with greater mitochondrial network aging rate also exhibit greater global transcriptional changes (including MMD, cell-cell variation, and gene variability). Nonetheless, there are cell types where different aspects of global aging patterns are inconsistent with each other. For instance, the ALA neuron (cluster 140_0) has a pronounced increase in cell-to-cell variation with age and a relatively small change in MMD. Thus, aging could be driven either by cell-type dependent conserved programs or in a more cell-type independent and chaotic way. A comparable effect exists during replicative aging of *S. cerevisiae* in which several aging events were observed, leading to distinguishable death phenotypes (Li et al., 2020). Finally, we note that in some cell types, cell-to-cell variability actually *decreased,* indicating that the paradigmatic, entropic, loss of order need not accompany aging.

In summary, we present in this study a near-comprehensive single-cell atlas for adult *C. elegans* gene expression with cell type-specific RNA signatures, allowing an unprecedented comparison of specific cell type aging. Gene expression only represents one layer of the overall cellular complexity, but the differential regulation we observe exposes a surprising aspect of cellular aging– a minor shared aging signature and an unanticipated level of divergence between the different cell types. The dysregulation of energy metabolism, including mitochondrial respiration, is the most universal change; a decline in ER function is pervasive as well, yet to a lesser extent. The diversity of aging trajectories is remarkable because it involves essential functions like the ribosomal protein genes, the proteasome or DNA repair. It is particularly surprising considering the low complexity of the *C. elegans* lineage, whose somatic cells are all post-mitotic. Significantly, the age-related gene expression shifts are not random but occur at the level of entire functional gene sets, suggesting a dynamic reorganization in patterns of cellular homeostasis. Another layer of dysregulation leads to increased cell-to-cell variability within cell types, one that is only modestly correlated with the functional changes in expression. In addition, our dataset allows the study of TF expression and activity changes with age, leading to the unexpected discovery that the expression of many TFs that promote longevity increases animal-wide with age. We believe this atlas will help the community to analyze and interpret the aging process with increasingly high levels of resolution, leading to a better understanding of how individual cells and tissues change their patterns of gene expression in ways that may promote, or forestall, a global loss of homeostasis and resiliency.

### Contact information for resources

Further information and requests for resources and reagents should be directed to and will be fulfilled by the lead contacts, David Kelley and Cynthia Kenyon. *C. elegans* strains generated in this study are made available upon request. Strains will be made publicly available through the Caenorhabditis Genetics Center (CGC) after the first personal request. Any additional information required to reanalyze the data reported in this paper is available from the lead contacts upon request. Single-cell RNA sequencing data have been deposited on our website at <u>c.elegans.aging.atlas.research.calicolabs.com/</u> as well as on GEO (link available upon request) and are publicly available as of the date of publication. Accession numbers are listed in the table S15.

## Methods

### Experimental model and growth conditions

All *C. elegans* and bacteria strains as well as the products used in this study are detailed in Table S15.

Worms were cultured on Nematode Growth Media (NGM) following standard methods, at 20°C or 25°C as indicated in OP50 *E. coli* (Brenner, 1974). Live *E. coli* colonize old worms leading to their premature death (Podshivalova et al., 2017). We chose to feed our population with killed OP50 to collect data during the geriatric period of the end of their life. For dead OP50, bacteria were treated with antibiotics, the only method we found to efficiently kill *E. coli* without altering their nutritive property. OP50 was grown in LB at 37°C overnight to saturation, then diluted in a large volume to OD 0.2 and cultured at 37°C until it reached OD 1 to 2. Gentamicin sulfate was then added to the culture at 200µg/mL and kept in the incubator for 5 hours. The resulting culture was centrifuged to pellet bacteria and resuspended in 1/75 the initial volume of LB gentamicin sulfate 16.7µg/mL.

2-layer agar plates were used to prevent worms from digging and to protect them from drying out. 15 mL autoclaved regular NGM containing 2% agar was poured in 10 cm plates. After 24 hours, 25 mL autoclaved NGM with 3% pure agarose was poured on top. Both layers contained gentamicin sulfate 16.7 µg/ml and carbenicillin 25 µg/ml, added in the solutions after autoclaving. After 24 hours, we added 0.5 mL dead bacteria solution under a sterile hood and we added another 0.5 ml when the lawn was dry. The plates were stored at 4°C.

For an aging time course assay, a population of approximately 200,000 *C. elegans* was required. Twenty *gon-2(q388)* L4s larvae (P0) were sampled and transferred to 5 regular OP50 6cm plates (twenty per plate). The F1 eggs were collected in S-Basal-PEG without bleach according to (Roux et al., 2016) and transferred to new plates. 3 days later, the F2 eggs were collected from plates without males, and 300-500 of them were added to new plates. At the end of day-1 of adulthood, the F2s lay thousands of eggs per plate (we added a few drops of 50x concentrated OP50 to avoid starvation). At least 30,000 F2s eggs were collected in solution using the same method, they were used as parents of the last generation. They were washed 4 times in 12 mL S-basal-PEG and left overnight at 20°C to hatch. 2500 F2 L1s were transferred on each 10 10cm OP50 plate (25,000 L1s total). They were incubated 48 hours at 20°C until they reached the L4 stage. 300 µL of 50x concentrated OP50 were added on top of the bacterial loan to avoid starvation. They were then shifted to 25°C and incubated 24 hours. A total of 300,000-400,000 eggs were recovered. Parents were discarded by centrifugation and the eggs were treated for 30 seconds in bleaching solution, washed immediately 5 times in S-Basal-PEG and incubated in solution at 25°C for 12 hours to allow hatching (100% of them should hatch). We estimated the L1s concentration using a stereo-microscope. We transferred around 200,000 L1s into ‘2-layers’ 10cm-plate with 2500 L1s per plate. 0.5 mL of dead OP50 was added to the plates every day from day 4 to day 10. We closely monitored the plates to discard the contaminated ones. Most worms were gonad-less, but because *q388* mutation in *gon-2* is not fully penetrant at 25°C, some worms retained their fertility (about 1-2 eggs per 100 worms). For this reason, we kept the population at 25°C the entire aging time course to prevent the development of the fertile worms. Spontaneous apparition of rare males is tolerated, they represented around 1% of the last generation.

For the same experiment in live bacteria, we replaced dead bacteria with live 50-times concentrated saturated OP50 and used 2-layers 10cm plates without antibiotics.

The two strains CF4596 and CF4569 were grown in presence of 1µM auxin passed the L4 stage of their final generation. Both strains contain an AID tag in C-terminus of *daf-2* locus, included for potential follow up studies under *daf-2(-)* condition. The rationale for using CF4596 strain in this study over an intact *daf-2* strain is that it will serve in future studies as a control against the long-lived strain CF4569 grown with auxin. We confirmed previous observation (Venz et al., 2021) that the AID insertion in the *daf-2* C-terminal locus did not impair DAF-2 function in absence of TIR1 or auxin (absence of abnormal dauer formation at 27°C, data not shown). The control was the same strain without the AID (CF4649, see below). CF4596 does not express TIR1 and is insensitive to auxin. CF4569 expresses TIR1 that can target DAF-2::AID to degradation upon auxin treatment (Zhang et al., 2015). This strain behaves like a *daf-2* loss-of-function in presence of auxin (100% dauer and long lived, not shown and (Venz et al., 2021)). During our preliminary experiments, CF4569 as well as a population of CF4086 grown in live OP50, were single-cell sequenced (see *Single cell gene expression quantification* paragraph below for details). We used these datasets to help increase the resolution of the high-dimensional single-cell data plotting (UMAP) for the condition we analyzed in depth, CF4596 in killed bacteria.

Our use of a mutation like *gon-2(q388)*, which prevents somatic gonad and germ cell development, was required to remove the overabundance of germ cells that mask the somatic cell-signal. There were two potential concerns with this strain. First, the gonad is required for development of certain other organs such as the vulval. However, since the strain is not fully penetrant, we were still able to identify gonadal and gonad-dependent cells though presumably at a relatively low abundance (see below, “Comparison to CF512 *gon-2^+/+^* strain)”. Second, loss of the germ cells is known to extend worm lifespan through activation of multiple stress-resistant transcription factors. However, this life extension requires the presence of the somatic gonad (Hsin and Kenyon, 1999), which is missing in *gon-2* mutants. As expected from laser ablation studies (Hsin and Kenyon, 1999), *gon-2(q388)* mutation has only a marginal effect on lifespan (David et al., 2010).

### Strain editing

Nucleotide sequences used for our CRISPR editing strategies are detailed in Table S15. For CF4596 and CF4569, *mu465[daf-2::AID]* allele was generated by inserting AID in the C-terminal of the *daf-2* locus. A CAS9-nuclease directed endonuclease reaction coupled to an oligo of the AID flanked with two 31bp homologies covering the outside region of *daf-2* stop codon was used to direct the homology-based repair as described previously (Paix et al., 2015). As a selection method, we used dauer formation on 100 µM auxin. A more detailed protocol of our strategy is available upon request.

CF4649 was generated to be used as a control to test whether *mu465[daf-2::AID]* could affect DAF-2 activity (in absence of TIR1). CF4649 is a crispR edition of CF4596, where a double Cas9 cut of the AID was done to reverse *daf-2::AID* into *daf-2+/+* by homologous recombination using an oligo.

All our CrispR reactions were carried out with a *dpy-10(cn64)* heterozygous co-edition marker (Paix et al., 2015). All our final clones were selected to be *dpy-10*+/+.

### Single cell isolation

The following protocol is inspired by the study from Zhang and collaborators (Zhang et al., 2011) and from Kalestky et al. (Kaletsky et al., 2016). We implemented a list of modifications to apply the protocol to adults. Bulk RNA sequencing performed before and after the dissociation protocol during our preliminary tests has shown some marginal ER-UPR stress response induction due to the worm dissociation. Following a previous report (Kaletsky et al., 2016), we tried to add the RNA transcription inhibitor actinomycin D to the dissociation buffer to prevent stress response. None of the concentrations we tested prevented this response. Instead, we found that shortening the time between the SDS/DTT treatment and the mechanical extraction on ice was the best way to limit this response to insignificant levels.

Approximately 30,000 CF4596 worms were harvested per time point and washed 4 times in S-Basal/PEG 0.01%. During the last wash, we harvested 1500 worms that we snap-froze in 1mL trizol to extract total RNAs to control for the cell extraction related stress. The other worms were pelleted in a 15mL tube. From the SDS/DTT treatment and before the cells were isolated in solution at 4°C, the procedure was conducted as fast as possible within 20 to 25 minutes. We measured the volume of the worm pellet (around 500µL) and added 3 times the volume of lysis buffer to it (200 mM DTT, 0.25% SDS, 20 mM HEPES pH 8.0, 3% sucrose, stored at –20°C and used freshly thawed). The time of incubation in the lysis buffer was variable and critical. It was carefully monitored under a stereo-microscope using 5 µL aliquots. We stopped the reaction before the animals’ cuticles broke but when the cuticles showed signs of loosening. In young worms, the head started to swell, and the mouth was starting to protrude. In old worms, this time was shortened by 1-2 minutes. Old worms were ready when the vulva of some of them started releasing internal substance. The animals’ body thrashing slowed down but they remained alive. The incubation time varied between 3 to 6 minutes. We stopped the reaction by adding 12 mL of M9/PEG, 0.01%. The worms were quickly pelleted at around 200 rcf for 30 seconds. We repeated this wash twice and resuspended the pellet in 3 mL M9/PEG 0.03% to transfer it into a 5 ml eppendorf tube. After pelleting the worms at 150 rcf for 20 seconds, the supernatant was removed and replaced with 1 mL freshly prepared pronase solution in L15 Leibovitz medium (L1518 Sigma adjusted to 340 m Osm with sucrose 28mM final; pronase 15mg/mL) as described in (Zhang et al., 2011). The pronase incubation was carried out at room temperature with gentle rocking for 5 to 15 minutes until about 5-30% of the worms broke open. The tube was then placed on ice while the solution was passed through a 21 G 1-inch needle in a 1 mL syringe back and forward 30 times. If necessary, a few additional passages through a 25 G 1-inch syringe were added. The release of cells gave the solution a cloudy aspect. The debris and the unopened worms were separated from the cells at 150 rcf for 1 minute and the supernatant (cell suspension #1) was transferred to a new tube. 250uL L15-FBS 10% (FBS heat inactivated) was added to the 1 mL cell suspension #1 at 4°C to inactivate the pronase. The debris pellet was then treated again with 1 mL fresh pronase solution rocking at room temperature for an additional 3 minutes and returned on ice for a second mechanical cell extraction. The solution was passed through the 21 G needle another 30 times (cell suspension #2) and 250 µL L15-inactivated FBS 10% was added to it. The two cell suspensions were then pooled at 4°C and passed through a 35 µm nylon mesh (C. Murphy lab protocol), pre-wet with L15 340mOsm 2% inactivated FBS, to filter debris. We used a commercial cell strainer as the nylon mesh holder, after its original filter was cut out (Suppl. Notes 3A). The resulting cell suspension was centrifuged 7.5 minutes at 500 rcf 4°C. The resulting pellet was gently suspended in 3 mL 4°C egg buffer BSA (8 mM NaCl, 3 mM MgCl_2_-6H_2_O, 3 mM CaCl_2_, 5 mM HEPES, 48 mM KCl, pH 7.3, BSA added the same day 10 mg/mL). Hemocytometer C-Chip was used to estimate the cell concentration through a DIC microscope (see below), before and after the cell sorting. First, the cell suspension was centrifuged 7.5 minutes at 500 rcf 4°C and the pellet was gently resuspended in 400 µL 4°C egg buffer with 10 mg/ml BSA. Four mL 4°C methanol 100% was added very slowly to the cells while slowly vortexing and the fixed cells were stored at −20°C. In the hemocytometer, small and larger cells could be seen (Suppl. Notes 3B). The cell count was between a few million up to 20 million cells in total per isolation. If the cell count was too low, the SDS/DTT incubation time was increased.

### Cell sorting and single cell RNA sequencing

It was always preferable to use a minimum number of 3-4 million cells maintained at 4°C. We found that the use of re-hydrated methanol-fixed cells was the best way to preserve cellular RNA integrity and was compatible with the 10x chromium droplet encapsulation technology. The cell suspension in methanol was centrifuged 7.5 minutes at 500 rcf 4°C and the cells were gently re-hydrated with 2 mL egg buffer BSA with 2U/µL RNase inhibitor. DAPI was added at 3 µM and incubated for 10 minutes on ice. The cells were filtered through a 40 µM strainer and sorted with BDFACS Aria Fusion with the 100µm nozzle running at a frequency of 31.0 kHz. Cytometer performance was assessed by running CS&T QC beads. Sort drop delay was determined using Accudrop beads. DAPI was excited with the 405 nm violet laser and its emission detected with a 450/50 bandpass filter. DAPI positive cells were selected by plotting Forward Scatter (FSC) vs. DAPI-compatible filter and gating on events positive for DAPI, selecting against negative events which we presumed to be debris. Among DAPI positive events, only the 2N DNA population was gated (Suppl. Notes 3C) and sorted into 1mL PBS BSA 2x RNAse inhibitor 4U/µl. The use of cells from early day 1 adult cells was necessary to spot the 1N, 2N and 4N DNA cell populations, older worms lost the 1N DNA population. A minimum of 50,000-80,000 cells were sorted to load a single 10x Chromium channel. The sorting solution was transferred to a 5 mL eppendorf and centrifuged 7.5 minutes at 500 rcf 4°C, the pellet was resuspended in PBS BSA 10 mg/ml with 2U/µl RNAse inhibitor. The flow-sorted somatic cells were processed into libraries using 10x Genomics Chromium Single Cell 3’ V3 Chemistry as directed by the manufacturer, with a few exceptions.

16,000 cells were loaded to aim for 10,000 cells per channel processed into a 10X Genomics Chromium chip, 3 channels per time point were used. cDNA was amplified for 12 cycles. Between 1/4 and 7/8 of the cDNA was moved forward into library construction, and the index PCR was cycled 12 times. Final libraries were brought to 2.4 nM and processed on an Illumina cBot and HiSeq4000 sequencer.

### Healthspan

Healthspan was measured as the decline of movement over time at 25°C with live OP50 or HT115 *E. coli*. It was recorded using an automated version of the Multi-Worm Tracker (MWT) computer vision system (Kerr et al. 2022), with images acquired using PixeLINK PL-D725MU cameras (5 megapixels, configured to run at 50 frames per second) with Navitar NMV-25M1 lenses (25 mm focal length, F 1.4 aperture). Worms were segmented using the core Multi-Worm Tracker algorithms available from https://github.com/ichoran/mwt-core.git, and analyzed using Choreography from https://github.com/ichoran/choreography.git (Swierczek et al., 2011). The MWT was run using a target object size of 12 to 200 pixels and using the new asymmetric background adaptation rate feature to be more sensitive to slow-moving animals (rate 5, asymmetry −3, for 8-fold faster acceptance of new dark regions than removal of old dark regions). Choreography was run using −p 0.04 −M 1.5 −t 30 −s 0.2 −S –shadowless options and the Reoutline::exp, Respine, and SpinesForward standard plugins, and −N all -o area,speed,midline,loc_x,loc_y was used to output a time course for each animal tracked. With these parameters, an animal had to have moved at least half a body length anytime within a 1 min time interval to be detected, but could be picked up in as little as 5 seconds if it moved enough. After detection, tracking continued even if the animal was still. The Choreography output was analyzed using custom code to calculate maximum speed (computed as displacement over 0.2 seconds, then median filtered using a 5-sample window) over various time intervals; this code is available at https://github.com/ichoran/metrology.git. The animals were assayed as follows: prior to recording, three mechanical taps were delivered to the animals’ housing to induce movement. Recordings started within 5 seconds and lasted for 490 s: animals were first allowed to calm down from being handled (300 s), then subjected to twelve taps at a 10 second inter-stimulus interval. Animals respond to this tap train with a prolonged increase in speed, in addition to exhibiting per-tap responses; each animal’s maximum speed was measured 30-40 seconds after taps ceased. Tracking was performed using 4 separate plates per condition, starting with approximately 40 L4 animals per plate. Animals were tracked every 6 hours for their whole lifespan. We computed the average maximum velocity of each worm population (plate) because of all movements measures, maximum velocity correlates the best with longevity according to (Hahm et al., 2015). 0 on the Y axis corresponds to the late L4 stage. A publication dedicated to the high-throughput automated multi worm tracker is in preparation (Kerr et al. 2022).

### Lifespan

All lifespans assays were carried out at 25°C with live OP50 or HT115 *E. coli*. Lifespans were measured either by the standard method (referred to as Manual) or using consecutives images from the Multi-Worm Tracker (referred to as Automated). Manual lifespan was scored by testing movement as described previously (Apfeld and Kenyon, 1999). Animals were scored as dead when they failed to move after prodding. Automated lifespan was scored by looking at a series of pictures of the same plate using ImageJ. Consecutive images of the same plate are captured every 6 hours as part of the healthspan assay; these were overlaid in red, green, and blue to provide a visual cue of movement: motionless animals are in shades of gray, while even animals that only move their heads slightly appear at least partly colored. Animals were scored as dead when they did not show any movement for a period of 18 hours. The lifespan curves shown in Figure 7 are representative of at least 3 independent experiments. All results are reported in Table S15. Lifespans measured in parallel manually or through automated imaging provided the same results (Rex Kerr et al, 2022).

### Candidate TF gene RNA interference screen for healthspan effect

To choose our candidates, we considered the universality and the extent to which TF gene expression or TF targets activity changed with age in clusters. Candidates were censored if they were known aging regulators in the GenAge database. We were limited to the RNAi clones that were present in our library (*C.elegans* Ahringer RNAi library from Source Bioscience Ltd)(Kamath et al., 2003) . We selected TFs based on the following criteria: (1) Highest number of clusters in which the TFs are expressed and significantly changing with age (15 TFs up-regulated, 11 TFs down-regulated). (2) Highest average fold change with age regardless of the cluster number (10 up-regulated, 3 down-regulated). (3) Most changing AUcell predicted TF targets activity changes (8 TF up-regulated, 8 down-regulated) (Table S15). RNAi feeding plates were prepared as follows. 750 uL of saturated bacterial culture was diluted with 750 uL LB 100 μg/ml carbenicillin and after 2 hours at 37°C, isopropyl β-d-1-thiogalactopyranoside (IPTG) was added at 4 mM concentration. After 4h at 30°C,150uL of the RNAi culture was added on a NG 1mM IPTG 100μg/ml carbenicillin plate. The bacteria were spread equally with a sterilized glass stick on the surface without reaching the plastic edges to reduce the thickness of the bacterial loan. The plates were incubated for 24h at 30°C. CF512 worms were grown for 2 generations at 15°C and harvested as overnight synchronized arrested L1s at 25C (see growth conditions for details). They were incubated 24 h at 25°C on OP50 NG plates to reach the L4 stage. For the initial healthspan screen, 40 L4s were added to each RNAi plate. The plates were sealed with parafilm with 4 holes to enable some gas exchange and their inside lid surface was treated with fogtech-DX anti-fog solution. Four plates per condition were loaded into the healthspan automated reader on separate racks, each rack containing one or two control plates. Our RNAi control strain was transformed with L4440 empty-vector plasmid. In the first batch, we screened 58 RNAi knockdowns, described in Table S15. In the second batch, we analyzed 11 RNAi if they showed some difference with control in the first batch. In the third batch, we analyzed the 5 confirmed RNAi hits, all of their previously observed movement phenotypes were confirmed (Table S15, S16). In parallel, manual lifespan measurements were performed on the best hits for the batch 2 and 3. The results are summarized in (Table S16). Additionally, we applied the RNAi treatments, corresponding to all five TF hits, starting from L1 larvae and followed their development. We found that none of the RNAis prevented or slowed down their growth, showing that the gene knockdown used were not essential or toxic to the worms.

### Microscopy

The worms were mounted between glass slides on fresh 4%-agarose pads in 1.5 uL 5 mM levimasole in S-Basal. The confocal microscope used for experiments in figures 1 and 6 was a Nikon Ti-E Microscope/Yokogawa CSU-22 Spinning Disk with a Photometrics Evolve camera, a Prior motorized stage with Piezo Z-drive, a NEOS AOTF adapter and controller for Micro-Manager 1.4 software. The objective lens was a Plan ApoVC 60X/1.4 with oil. The laser wavelengths used were 405, 491, 561, 640 nm at 100 mW. Images were analyzed with NIS Elements 4.13 and ImageJ/FIJI. Images shown are a maximum density projection of Z-stacks acquisition of the entire cell of interest with 0.3 μm steps.

Transmitted DIC microscopy images were captured using a Leica DM6B with a Micropublisher 6 digital camera (QImaging) and 40x 0.8 Dry FLUO/BF/POL/DIC objective. Images were acquired using µManager software. The dissecting microscope used was a Leica MZ16FA microscope (Leica, Bannockburn, IL, USA).

### Mitochondria morphology scoring

Worms were mounted individually as described above. Z-stacks of the cells of interest were acquired and maximum density projections were computed on NIS Elements 4.13. DIC images were used as a validation that the right cells were captured. Then, GFP fluorescence images were isolated and treated with photoshop to mask any background fluorescence or details that could inform on the age of the worms. The extent of mitochondrial fragmentation was scored blind by two persons independently as shown in Figure S6. If more than one cell of a given cell type was scored in the same animal, the scores were averaged. Every visible cell was imaged. The cells far from the objective were often blurry and impossible to score. If one of the two cell types could not be imaged, the worm was discarded. The worms imaged came from two independently aged populations.

### Single cell gene expression quantification

Single cell-sequencing libraries were prepared from three independent populations across multiple time points. A population of CF4086 was grown in live OP50 bacteria and harvested at days 2, 4, 6, 8, and 10 of adulthood. A population of CF4596 was grown in killed OP50 bacteria and harvested at days 1, 3, 5, 8, 11, and 15. A population of CF4569 was grown in killed OP50 bacteria and harvested at days 1, 3, 5, 8, 11, and 15. A total of 200,000 cells were sequenced. In the current study we focused only on the population grown in killed OP50 bacteria. This choice is explained in the “Experimental model and growth conditions” section. Additional cells were only used to learn cell embeddings and clustering in order to better perform cell type annotation.

Sequence alignment and counting were performed against *C. elegans* genome assembly CE11 WS268, with Cell Ranger 3.0.0, employing STAR alignment tool (Dobin et al., 2013). Following alignment, we implemented the approach in Cell Ranger to filter out empty droplets and used Cell Ranger output “filtered_feature_bc_matrices’’ for downstream analysis.

### Raw data process and initial filtering

We applied CellBender v2.1 on raw 10x feature barcode matrices for ambient RNA removal and empty droplet detection with parameters --expected-cells 10000 and --total-droplets-included 20000 (Fleming et al., 2019). The filtered matrices were then processed with the Scanpy single cell analysis toolkit for raw data quality control (Wolf et al., 2018). We filtered out genes expressed in less than or equal to 5 cells and filtered out cells with 20% or more UMIs from mitochondria or ribosomal RNA.

### scVI denoising with and without age-correction

We applied scVI 0.6.0 to learn latent representations of each cell and also to construct a denoised expression matrix (Lopez et al., 2018). We trained an scVI model with a single hidden layer of size 768, a latent space of 128 dimensions, and a dropout rate of 0.2 on all cells. The initial learning rate is set at 0.001 and early stopping was implemented with a patience of 40 epochs, lr_patience=20 and learning rate dropping factor of 0.1.

In addition to the scVI model described above, to annotate cell types *de novo* without age as a confounder, we re-preprocessed the data with scVI, controlling for the time point sample as a batch effect and decoding the latent embeddings back to gene expression space as denoised profiles (referred to as age-corrected model).

Subsequently, we used this age-corrected scVI model to perform clustering and annotation since we did not want changes due to aging to confound cell clustering and cell type annotation.

### Germline, sperm, embryonic cell annotation and removal

We curated marker gene sets of 136 genes for germline, 18 genes for sperm and 16 genes for embryonic cells (listed in Table S17). We implemented a modified version of AUCell to annotate germline, sperm and embryonic cells (Aibar et al., 2017).

For each of the marker gene sets, we computed an AUCell score for every cell using the top 10,000 highly expressed genes according to the scVI denoised expression matrix. Then we computed AUCell scores for every cell using 10,000 random size-matched gene sets. For each cell, we compared AUCell scores of target gene sets versus random gene sets to derive an empirical p-value quantifying the deviation from a null distribution. For sperm and embryonic gene sets, we annotated the cell to be sperm or embryonic if the FDR adjusted p-value was less than 0.01.

This thresholding strategy was insufficient for germline annotation, because the overall distribution of germline scores was shifted to the right of the null distribution. Since we observed that the AUCell scores for the germline gene set were clearly bimodal, we used Otsu’s threshold for germline annotation.

We detected 0.2% sperm cells, 0.3% embryonic cells and 11.0% germline cells in the dataset. These cells were removed computationally from downstream analysis (Suppl. Notes 1A).

### Doublet detection and removal

We applied Solo for doublet detection (Bernstein et al., 2019). Solo is trained to predict doublets for each library independently on top of the trained scVI representation, with cl_hidden = 128 and cl_layers = 1. We annotated 11.8% doublets in the live bacteria samples and 11.9% in the dead bacteria samples. These cells were removed from downstream analysis.

### Additional ambient RNA removal

We still observed ambient RNA contamination in the day-1 sample even after CellBender ambient removal (Suppl. Notes 1B). Among the most abundant mRNAs in the ambient profile (aggregated profile across empty droplets), we found cuticle collagen genes (including *col-140*, *col-119,* etc.) in almost all cells before ambient RNA and doublet removal. Even though CellBender successfully cleaned these cuticle collagen gene mRNAs in the majority of day-1 cells, there were still a number of cells in each cell-type cluster where it was not fully removed. Thus, we performed additional day-1 cell filtering to account for the remaining ambient RNA issue. For cells in day-1 animals, we first determined whether each cell type cluster had true cuticle collagen expression by examining the median *col-140* expression level. *col-140* is the most highly expressed collagen mRNA that we observed, and expression of collagen genes were highly correlated. Thus, we made the assumption that if *col-140* has 0 count in more than half of the cells for a cell type cluster, the cell type cluster does not express cuticle collagen genes and all the collagen mRNA that we observe in this cell type cluster is due to ambient RNA contamination. There were 14 clusters where collagen gene expression was real according to this criteria (which we call “cuticle_clusters”), the rest are “non-cuticle clusters”. The majority of them have already been described to express collagen: hypodermis, seam, CEPsheath, etc. For every cell in day-1 animals, we computed the AUCell score of GO term “structural constituent of cuticle”, which contains 220 collagen genes. We examined the distribution of the cuticle AUCell scores for “cuticle clusters” and “non-cuticle clusters’, and observed a clear separation. We removed all day-1 cells in the “non-cuticle cluster” with cuticle AUCell score >0.1. We removed a total of 4,698 out of 19,246 cells from the day-1 sample (Suppl. Notes 1B).

### Clustering and sub-clustering

Leiden algorithm was performed with default resolution of 1 on the nearest neighbor graph computed on scVI age-corrected latent representations (Traag et al., 2019). This resulted in 147 clusters we named *super-clusters*. We performed systematic and manual annotation to define an anatomy (cell type) associated with each super-cluster (see below for detail).

During super-cluster cell type annotation, we observed that some super-clusters have clear substructure with diverse annotations and aging trajectories. This suggested sub-clustering effort is required for accurate anatomy annotation.

For cells in each super-cluster, we re-computed a nearest neighbor graph based on the age-corrected latent representation, and performed community detection using Leiden with increasing resolutions: 0.01, 0.05, 0.1, 0.2, 0.3, 0.4, 0.5, 0.6, 0.7, 0.8, 0.9, 1. At each resolution, we visualized the sub-cluster structure on the uncorrected UMAP and compared that with the distribution of time points and anatomy AUCell scores. We then manually selected the resolution which allows cells from all time points to be represented within each sub-cluster while separating clearly different anatomy terms. For example, we determined that Leiden sub-clustering with a resolution of 0.6 was optimal for super-cluster 3, which resulted in 6 sub-clusters. This is the maximum resolution we could achieve such that most time points (ages) are represented in each sub-cluster (Figure 1B, S1E). Using this pipeline, after dividing some of the super-clusters, we obtained a total of 211 clusters.

### Cell-level AUCell anatomy scoring

To obtain a database of genes with known anatomic expression, we downloaded all available “Expression Pattern’’ gene annotations from WormBase (version WS269) using WormMine. “Expression Pattern’’ annotations are derived using expression reporters (e.g. GFP) or by performing *in situ* mRNA hybridization. For each anatomy, we only used the marker genes scored as “certain” by WormBase. The resulting matrix linked expression of 5375 unique genes to 2438 unique anatomical terms (WBbt IDs) that represented functional systems (e.g. nervous system), anatomies (e.g. head ganglion) or cells (e.g. ASIL, ABplaapapppa). We filtered this matrix to contain only cell-level anatomies. This resulted in a matrix linking 1950 genes to 532 cell types, with a median of 17 unique genes per cell anatomy (Table S1).

We used gene sets associated with these cell anatomies to compute anatomy AUCell scores for each anatomy term (Table S1), which we subsequently used for cell type annotation. For each of the anatomy terms, we computed AUCell scores for each cell using the top 10,000 expressed genes based on the age-corrected scVI denoised expression matrix.

### Anatomy annotation for each cluster

We applied two systematic and one manual annotation method to the 211 clusters for anatomy (cell type) annotation (Table S1). For cell type annotation, we used the age-corrected scVI denoised expression matrix.

#### Anatomy annotation based on differentially expressed genes (DEGs)

We first performed one-versus-all differential expression analysis to identify up to 200 DEGs for each cluster based on a Wilcoxon rank sum test on the denoised gene expression matrix (q-value < 0.05, log2FC > 0). Then for each DEG gene set, we performed a hypergeometric test against each of the 532 WormBase anatomy terms. After correcting for multiple hypothesis testing, anatomy terms with Benjamini-Hochberg adjusted p-value <0.1 were considered potential candidate anatomy associated with the super-cluster.

#### Anatomy annotation based on AUCell scores

AUCell was implemented to compute cell level anatomy scores as described above. In order to summarize the cell level AUCell anatomy scores at the cluster level, we computed a summarized score for each cluster by comparing the set of scores for cells in the cluster with scores of the rest of the cells using an AUC rank statistic analogous to the AUCell method. We then assigned each anatomy term to the top three clusters with the highest scores.

#### Manual annotation

We inspected the AUCell score distribution of each anatomy term on the entire UMAP and manually assigned each anatomy term to one or multiple Leiden clusters.

Finally, we compared the annotations for each cluster from the three methods. If different anatomies matched a given cluster, we picked the one with the highest average expression in this cluster. However, we prioritized the specificity over the level of expression; I.e. if one anatomy was more highly expressed than another one in a given cluster, the lower expressed one was retained if it was expressed solely in this cluster and the other one was not.

### Cluster-averaged transcriptome correlation

We computed aggregated cluster expression profiles by taking the average of all cells in each of the 211 clusters and projected the profiles to 50 dimensional space using PCA. Then we computed pairwise Pearson correlation for all clusters.

### Comparison to CF512 *gon-2^+/+^* strain

Because we used a strain carrying *gon-2(q388)* mutation that partially prevents the development of the gonadal tissues, we verified the integrity of these clusters by comparing them (strain CF4596) with those from CF512, a strain carrying a wild-type copy of *gon-2* that was profiled with single-cell RNA-seq for a preliminary experiment (full data set not published). CF512 carries *fer-15(b26)* and *fem-1(hc17)* alleles that prevent the normal development of their sperm cells.

We preprocessed the CF512 sample using the same workflow described above. 24,524 cells from CF512 cells passed quality control. We then performed Leiden clustering. We identified clusters associated with sheath cells, distal tip, vulva uv1 and uv3 in the CF512 sample using AUCell scores of the corresponding Wbbt anatomy terms (Wbbt:0005828, Wbbt:0004520, Wbbt:0006793, WBbt:0006791). We compared transcriptional profiles of cells in these clusters with the cells in the corresponding clusters in our CF4596 sample (Suppl. Note 1D).

### Comparison to Kaletsky et al. worm tissue bulk RNA-seq

We downloaded the tissue-specific marker genes for hypodermis, intestine, muscle and neurons from Table S9 of Kaletsky et al.’s publication (Kaletsky et al., 2018). We then computed the Jaccard index between their marker genes with marker genes for each of our 211 cell type clusters. Our marker genes are defined by absolute value of log2 fold change > 2 and adjusted p-value < 0.01 in the cell type differential expression analysis.

### Comparison to Taylor et al. L4 larvae sc-RNAseq

We downloaded the preprocessed neuron single cell expression matrix cds file from cengen.org (Taylor et al., 2021). We computed an average expression profile for each of the 115 neuron cell types from the Taylor et al. study and each of the 133 neuron clusters in our study. Then we computed the pairwise Pearson’s correlation between the transcriptional profiles of our neuron cell types and the neuron cell types annotated in Taylor et al. For each of the 115 neuron cell types in Taylor et al. study, we also reported the cell type cluster in our dataset that best correlates with it (Table S3).

### Comparison of intestinal cluster pre- and post-FACS

We processed the pre-FACS sample in the same way as the post-FACS sample. After filtering out germ cells in the pre-FACS sample, we performed Leiden clustering on the sample and identified the intestine cluster based on expression of key transcription factors *elt-2* and *elt-7*.

We evaluated the quality of the annotated intestinal clusters in the post-FACS sample by comparing to intestinal cells in the pre-FACS sample. We correlated all cell type clusters in the post-FACS sample with the intestine cluster in the pre-FACS sample. We also compared the overlap of marker genes derived from intestinal clusters from pre-versus post-FACS samples (Supp. Note 1E-J).

### Transcription factor activity inference

For motif analysis, we used the *C. elegans* motifs in the CIS-BP database for 269 unique TFs (Weirauch et al., 2014). We selected a single motif for each TF based on evidence precedence rules. First, we preferred directly-measured motifs over inferred motifs. In cases with multiple direct motifs, we chose the motif based on the following ranked order: ChIP-seq, HocoMoco, DeBoer11, PBM, SELEX, B1H, High-throughput Selex CAGE, PBM:CSA:DIP-chip, ChIP-chip, COMPILED, DNaseI footprinting.

We used the FIMO software to find motif hits for the 269 TFs in promoter regions of *C. elegans genes*, defined as −500bp upstream to +100bp downstream of the 5’ end in the WS271 gene annotation (Grant et al., 2011). If the gene falls in an operon, we considered the gene to be regulated by the promoter of the first 5’ gene in the operon. We ran FIMO with default parameters and defined significant hits at a 1e-4 p-value threshold. We constructed a binary TF-gene interaction matrix based on the presence of a significant FIMO motif hit in the promoter associated with the gene.

We defined the target gene set for each TF based on the binary TF-gene interaction matrix. We then inferred the TF activity for each cell by computing AUCell scores of the TF targets using the scVI denoised matrix.

In order to study the role of TFs in each tissue, for each TF-tissue combination, we computed the Spearman’s correlation between TF expression and TF activity (AUCell) across single cells after regressing out age as a covariate using scanpy.pp.regress_out. This is a conservative way to quantify TF-target relationship, since the only variation left in the data is cell-to-cell heterogeneity. We only considered TF-tissue combinations where there are more than 5 cells in the cell type cluster and the average expression of the TF is greater than 10 RPM. We then evaluated the significance of the correlation between expression and activity in each TF-tissue combination using a permutation test. For cells in a particular cell type cluster, we generated a null distribution of TF expression-activity correlations by sampling 10,000 fake expression-activity pairs and computing the Spearman’s correlations. A fake expression-activity pair is defined as a pair of expression and AUCell vectors such that the expression vector is from TF-A, and the AUCell vector is from TF-B and the motifs of TF-A and TF-B are sufficiently different from each other (Pearson’s correlation < 0.6). TF expression-activity correlations with a permutation test FDR adjusted p-value < 0.05 are considered significant.

### Differential expression analysis during between young and old cells

In order to characterize common and tissue-specific changes during aging, we performed differential expression analysis of young (d3, d5) versus old cells (d8, d11, d15) for each cluster using scanpy rank_genes_groups function for all genes using Wilcoxon test. A cell type cluster was skipped if there were less than 5 young cells or 5 old cells.

### Gene set enrichment analysis (GSEA)

For gene set enrichment analysis, we downloaded all GO pathways relevant for *C. elegans* using the R package EnrichmentBrowser, including Biological Process (BP), Cellular Components (CC) and Molecular Function (MF) pathways.

For each cell type cluster, we ranked all genes by the differential expression log2FC in the old versus young comparison. GSEA was performed on the ranked gene set using the command line tool gsea-v3.0 with 2,166 BP, CC and MF GO pathways after excluding any gene sets smaller than 5 or larger than 2000.

By default, GSEA tests for enrichment of each gene set in each cell type cluster in both directions (increasing and/or decreasing with age). We summarize the result by reporting the −log10 FDR q-value and normalized enrichment score for the direction which the gene set is more enriched in and assign a sign based on the direction (+ if enriched in genes that increase with age, - if enriched in genes that decrease with age).

To help with interpretation, we performed hierarchical clustering on the 2,166 GO pathways based on their normalized enrichment score across cell type clusters. Then we clustered the 2,166 GO pathways into 200 GO clusters and represented each GO cluster by the GO term with the largest number of genes.

### Dynamic gene expression changes

#### Gene trajectory analysis

In order to characterize nonlinear transcriptional changes during aging, we first grouped genes that share a similar aging trajectory into gene modules. For the 4,232 differentially expressed genes with a fold change > 2 and FDR-adjusted p-value < 0.01, we first computed a cell type-specific aging trajectory for each gene in each cell type cluster where it is expressed by z-scaling the expression vector across days 1-11. Day 15 was excluded from the trajectory dynamic gene expression analysis because there were too few cells. For each gene, we computed an averaged trajectory across cell types. We identified modules by hierarchical clustering on the cell-type-averaged trajectory with Euclidean distance and complete linkage and then cut the tree into six modules. We associated each gene module with GO pathways by performing GO enrichment analysis using R package “GOstats”. For each of the gene modules, we also computed a module aging trajectory for each cell type cluster by taking the average across genes.

#### Time-series GSEA analysis

We performed differential expression analysis for each cell type cluster for each pair of consecutive time points except for d15 (d1 vs. d3, d3 vs. d5, d5 vs. d8 and d8 vs. d11) using the scanpy rank_genes_groups function, similarly to the previous young versus old analysis. We performed GSEA using the command line tool gsea-v3.0 with 2,166 BP, CC and MF GO pathways on the genes ranked by the “score” column for each comparison. We considered a pathway to be significantly associated with aging if the FDR-adjusted p-value < 0.01.

### Maximum mean discrepancy

Following prior work, we implemented maximum mean discrepancy (MMD) to quantify the magnitude of transcriptional change during aging (Kimmel et al., 2020). MMD quantifies the difference between two populations p and q given samples from each population by taking the difference between the mean function values of the two samples over some smooth function f (Gretton et al., 2012). Intuitively, when the function f is rich enough, the difference between the mean function values will be 0 if and only if p=q. In our case, we apply a radial basis kernel function and the MMD between young cells and old cells for each cluster type cluster is computed as:

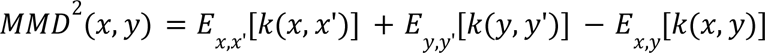

Where x represents sample from the young cell population, y represents sample from the old cell population and 𝑘 represents the radial basis kernel (Gretton et al., 2012).

### Cell-cell variation

We quantified the transcriptional variation for a group of cells (cell-cell variation) by measuring the Euclidean distance of each cell to the group centroid in gene expression space (Kimmel et al., 2019). We computed this statistic for cells in each cell type and each time point. Then for each cell type cluster, we estimated the rate of change in cell-cell variation per day. For a specific cluster, if the cell-cell variation at d1 is A and at d3 is B, then the rate of change per day is estimated by (B/A)^(½). We computed the rate of change for each pair of consecutive time points and reported the average across time points.

### Gene variability

We quantified the biological variability in mRNA abundance (gene variability) within a homogenous cell population (same age and cell type) by fitting a negative binomial regression model to each mRNA species and computing the variance of the analytical Pearson residual(Hafemeister and Satija, 2019; Lause et al., 2020). The Pearson residual is intuitively a measurement of goodness-of-fit with respect to a null distribution assuming all cells share the same transcriptional profile (Lause et al., 2020). To focus on variance at the cell-type level, we summed every gene’s statistic for each cell type cluster and generated a cell type by time point matrix of gene variability estimates. Then we computed the rate of change in gene variability with age for each cell type cluster similarly to the analysis we performed for cell-cell variation.

### Transcription factor aging analysis

For each of the 230 TFs in the CIS-BP database, we evaluated the change in TF expression during aging in each cell type cluster by computing the TF expression log2FC in young (d1, d3, d5) versus old (d8, d11, d15). We evaluated the changes in TF activity during aging by the difference in mean AUCell scores in young (d1, d3, d5) versus old (d8, d11, d15) cells.

### microRNA regulation during aging

We downloaded predicted targets for each of the 60 miR families in the targetScan database release 6.2. For each cell type cluster, we evaluated the enrichment of targets for each of the 60 microRNA in significantly up-regulated and significantly down-regulated genes (|log2FC|>1 and FDR-corrected p-value<0.01) using a hypergeometric test. We considered a microRNA target gene set to be significantly enriched in the age-associated up-regulated (or down-regulated) genes if the FDR corrected hypergeometric test p-value <0.01.

### DEXICA module analysis

GSEA was performed for each cell type cluster in young vs. old cells as described above, but using 418 DEXICA hemi-modules (Cary et al., 2020) as gene sets. Hemi-modules were ranked by the total number of cell type clusters with significant (FDR-adjusted p-value < 0.01) enrichment with age (in either direction) to identify the top 25 age-related DEXICA hemi-modules (detailed in Table S18). Hierarchical clustering was performed on cell type clusters but not hemi-modules, preserving hemi-module order from most to least frequently enriched with age. m36a comprises 487 genes and changes in expression of these genes (as calculated using AUCell scores) at each time point were visualized in significant cell-type clusters.

### Limitation of the study

Our isolation of *C. elegans* adult cells allows for a near complete coverage of cell types for sequencing. To do so, we used genetics as well as a ploidy-based sorting method in order to enrich somatic cells against a large amount of germ cells. As a result, our final dataset contains a small amount of germ cells that have been selected to be diploid and are not representative of the diversity of germline cells and oocytes with N and 4N ploidy.

The intestine is another cell type affected by the sorting step. Intestinal nuclei undergo endoreduplication during early development to become polyploid(Hedgecock and White, 1985). Nevertheless, our final UMAP contains several intestinal clusters. These clusters should be considered carefully. Our manuscript features an additional section with an independent dataset for this cell type derived from a prior sequencing experiment without ploidy sorting.

## Author’s contributions

AR designed and performed animal experiments and cell extraction, analyzed data, contributed intellectually and wrote the paper. HY designed and performed most scRNA-seq data analysis, contributed intellectually and wrote the paper. KP performed data analysis involving DEXICA gene expression modules, contributed intellectually and edited the paper. DH performed sequencing data acquisition and analysis, contributed intellectually and edited the paper. RK provided technology and guidance for the *C. elegans* Observatory. CK contributed to experimental design, data interpretation and the manuscript. DK performed data analysis, contributed intellectually and wrote the paper. The authors are members of the Kelley, Kenyon and Genomics labs at Calico Life Sciences LLC.

## Acknowledgements

We are grateful to many members of Calico Life Sciences LLC. We would like to thank particularly important contributions. Andrea Ireland, Twaritha Vijay and Margaret Roy assisted with single cell RNA-sequencing experiments. Maria Ingaramo and Andrew York assisted with our microscopy. Alex Chekholko and Adam Baker supported our needs for setting up the website. Ashok Shah and Gerard Balon ensured our supplies of media and worm plates. Jerome Goudeau, Peichuan Zhang and Madhuja Samaddar provided technical assistance related to our *C. elegans* experiments. Jonathan Paw helped with cytometry. Jacob Kimmel and Vikram Agarwal provided helpful advice for our data analysis. Outside Calico, strains from the C. elegans Genetics Center (CGC), which is funded by the NIH Office of Research Infrastructure Programs (P40 OD010440), were used during testing and development. Data collected by the WormBase website, http://www.wormbase.org, release WS284 (2021), were used in this study.

