## Supplemental files and notes for "The complete cell atlas of an aging multicellular organism"

**Supplemental figures**
Page 1-14

**and supplemental notes**
Page 15-26

### The complete cell atlas of an aging multicellular organism

Antoine E. Roux[1,2], Han Yuan[1,2], Katie Podshivalova[2], David Hendrickson[2], Rex Kerr[2], Cynthia Kenyon[2,*] and David R. Kelley [2,*]

[1] Authors contributed equally
[2] Calico Life Sciences LLC, South San Francisco, California 94080, USA

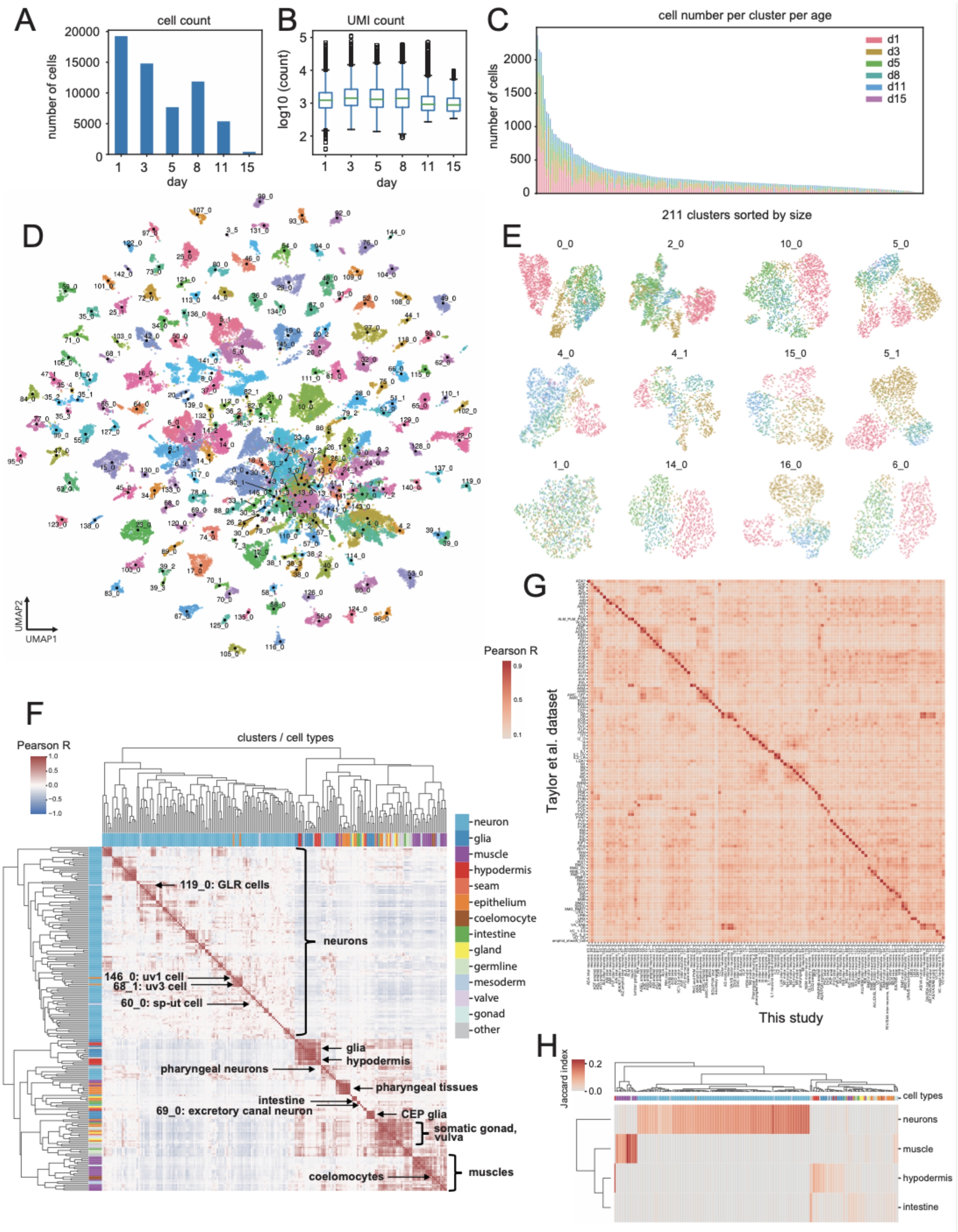

A
cell count
number of cells
20000
15000
10000
5000
0
1 3 5 8 11 15
day
B
UMI count
log10 (count)
5
4
3
2
1 3 5 8 11 15
day
C
cell number per cluster per age
d1
d3
d5
d8
d11
d15
number of cells
2000
1500
1000
500
0
211 clusters sorted by size
D
UMAP2
UMAP1
E
0_0
2_0
10_0
5_0
4_0
4_1
15_0
5_1
1_0
14_0
16_0
6_0
G
Pearson R
0.9
0.5
0.1
Taylor et al. dataset
This study
F
Pearson R
1.0
0.5
0.0
−0.5
−1.0
clusters / cell types
neuron
glia
muscle
hypodermis
seam
epithelium
coelomocyte
intestine
gland
germline
mesoderm
valve
gonad
other
119_0: GLR cells
neurons
146_0: uv1 cell
68_1: uv3 cell
60_0: sp-ut cell
glia
hypodermis
pharyngeal neurons
pharyngeal tissues
intestine
69_0: excretory canal neuron
CEP glia
somatic gonad, vulva
coelomocytes
muscles
H
Jaccard index
0.2
0.0
cell types
neurons
muscle
hypodermis
intestine

**Figure S1.** Raw sequencing data, clustering and cluster annotation of the dataset, related to Figure 1.

A) Bar plot showing the number of cells recovered at each time point. B) Box plot showing the distribution of log10 UMI counts for cells at each time point. C) Number of cells from each time point in each cluster. Clusters sorted by total cells in the cluster. D) UMAP representation for 211 annotated cell type clusters colored by cluster. E) UMAP visualization for top 12 largest cell type clusters. F) Heatmap of Pearson correlations between every pair of cell-type clusters' average gene expression. High-level tissue type is annotated in the color bar. Clusters of interest and tissues are labeled. G) We computed the cell type cluster-averaged gene expression profile for our neuron clusters and compared the profiles with cluster-averaged profiles from the Taylor et al. study using Pearson's correlation. We then compared ours and Taylor et al. annotations for the best matching clusters (Table S3). H) Heatmap of Jaccard indices between cell type specific markers derived from neurons, muscle, hypodermis and intestine from Kaletsky *et al.* study and cell type specific markers for each cell type cluster from our analysis (Kaletsky et al. 2018).

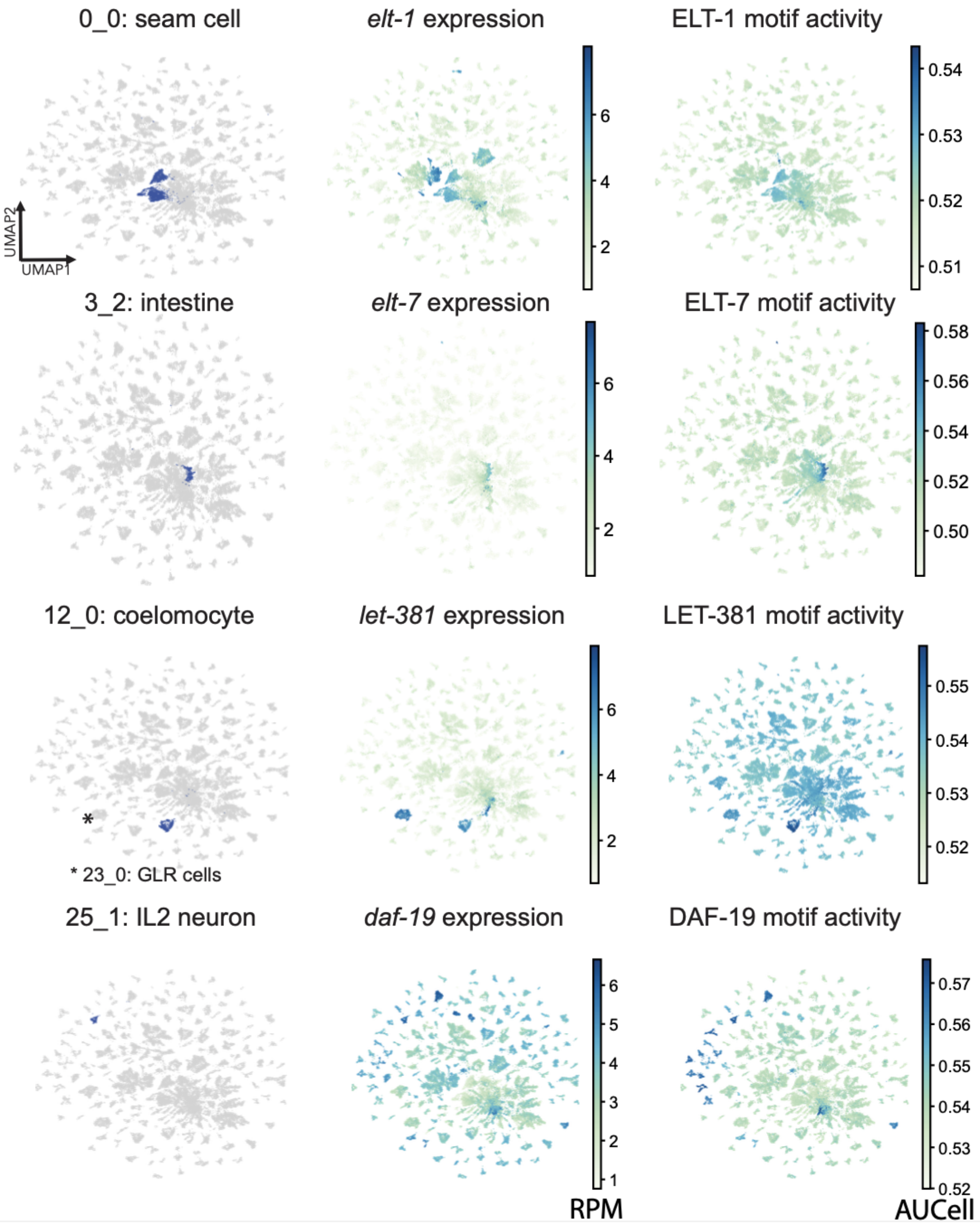


**Figure S2**. Examples of tissue-specific associations of the TF expression with its target genes activity (motif-based modules), related to Figure 2.

For each row, from left to right: (1) UMAP for all cells, with the cell type cluster of interest labeled in blue; (2) UMAP showing TF gene expression in log2(RPM); (3) UMAP showing AUCell scores for TF targets, as defined by motif hits; (4) UMAP showing AUCell scores for TF targets, as defined by the intersection of motif hits and co-expression. From top to bottom: seam cell specific activity of ELT-1; intestine specific activity of ELT-7; coelomocyte specific activity of LET-381; IL2 neuron specific activity of DAF-19. * locates the cluster 23_0, GLR cells.

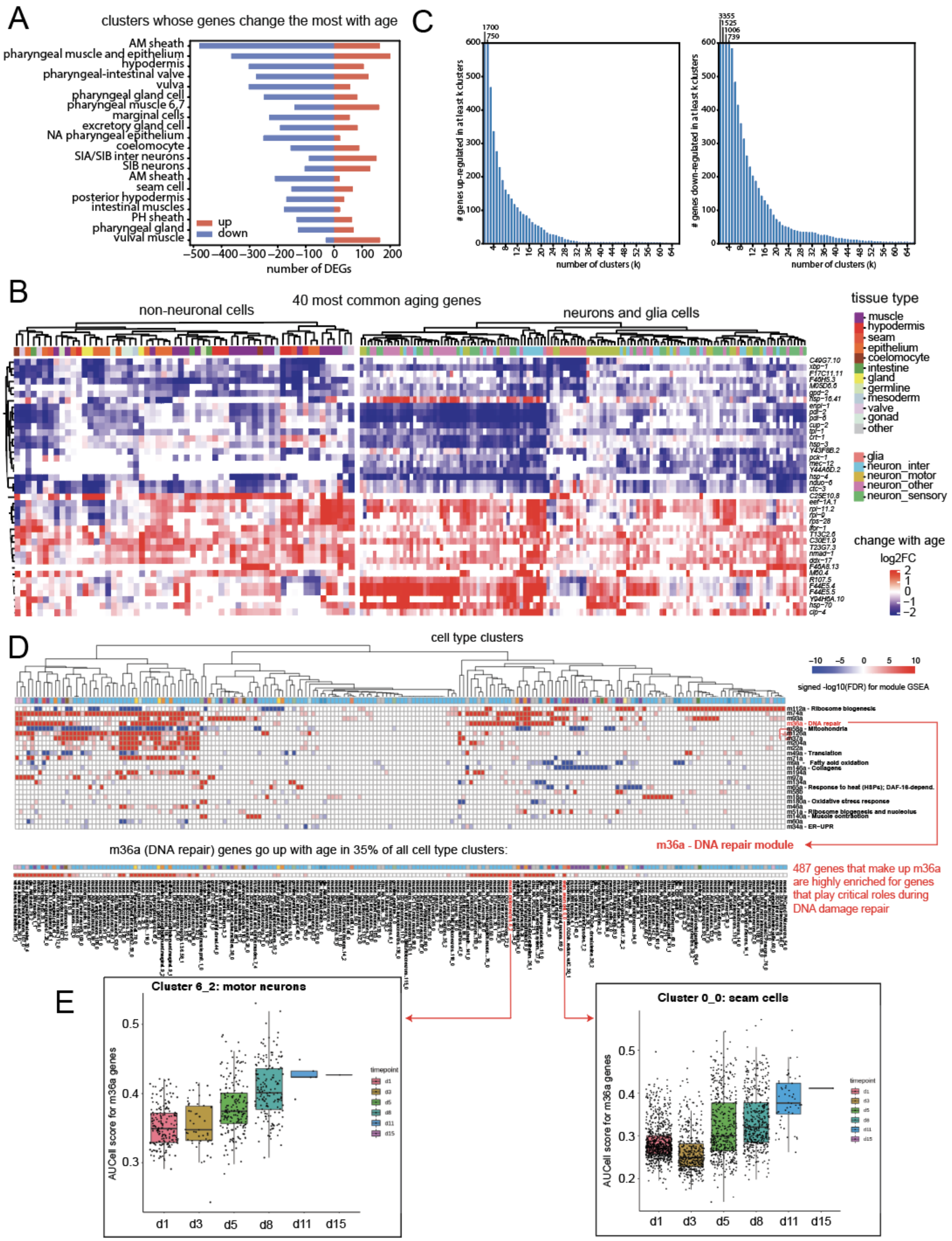

A
clusters whose genes change the most with age
AM sheath
pharyngeal muscle and epithelium
hypodermis
pharyngeal-intestinal valve
vulva
pharyngeal gland cell
pharyngeal muscle 6,7
marginal cells
excretory gland cell
NA pharyngeal epithelium
coelomocyte
SIA/SIB inter neurons
SIB neurons
AM sheath
seam cell
posterior hypodermis
intestinal muscles
PH sheath
pharyngeal gland
vulval muscle
up
down
number of DEGs
C
### genes up-regulated in at least k clusters
### genes down-regulated in at least k clusters
number of clusters (k)
B
40 most common aging genes
non-neuronal cells
neurons and glia cells
tissue type
muscle
hypodermis
seam
epithelium
coelomocyte
intestine
gland
germline
mesoderm
valve
gonad
other
glia
neuron_inter
neuron_motor
neuron_other
neuron_sensory
change with age
log2FC
D
cell type clusters
signed -log10(FDR) for module GSEA
m112a - Ribosome biogenesis
m36a - DNA repair
Mitochondria
m49a - Translation
Fatty acid oxidation
Collagens
Response to heat (HSPs); DAF-16-depend.
Oxidative stress response
Ribosome biogenesis and nucleolus
Muscle contraction
m34a - ER-UPR
m36a (DNA repair) genes go up with age in 35% of all cell type clusters:
m36a - DNA repair module
487 genes that make up m36a are highly enriched for genes that play critical roles during DNA damage repair
E
Cluster 6_2: motor neurons
Cluster 0_0: seam cells
AUCell score for m36a genes
timepoint
d1
d3
d5
d8
d11
d15

**Figure S3.** Cell-type specific aging signatures, related to Figure 3.

A) Top 20 clusters with the most number of genes that change with age. Blue bars show genes that change in the negative direction. Red bard show genes that change in the positive direction. B) Top 40 genes that are differentially expressed during aging across most cell types. Heatmap shows the log2 fold change between young and old for each gene in each cell type. C) left) distribution of genes with significantly increased expression in at least K clusters; right) distribution of genes with significantly decreased expression in at least K clusters. D) Top 25 DEXICA gene hemi-modules that change with age. GSEA analysis comparing gene $\log_2$ fold-changes between young (d1, d3 and d5) and old (d8, d11 and d15) cells in every cluster was conducted using 418 DEXICA hemi-modules as gene sets. High-confidence annotations are shown for those hemi-modules, where they are available. The lower panel highlights the result for m36a, which represents DNA damage repair. E) Left: Boxplot with jitter showing expression of m36a DEXICA hemi-module gene set in the motor neurons cluster 6_2 in log (count per 10k) at each time point; Right: boxplot with jitter showing expression of m36a DEXICA hemi-module gene set in the seam cells cluster 0_0 in log (count per 10k) at each time point.

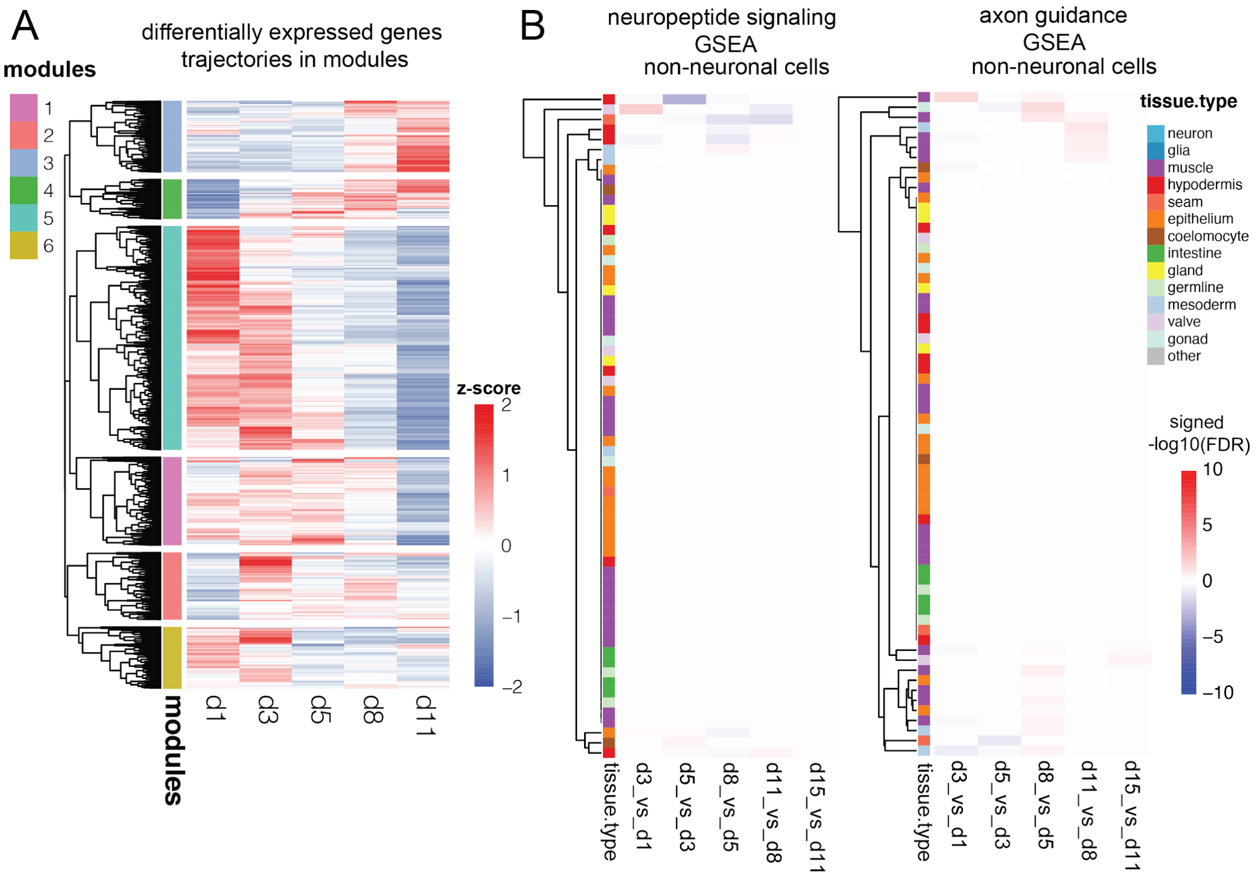


**Figure S4**. Differentially expressed genes grouped into six aging trajectories using hierarchical clustering, related to Figure 4.

A) Heatmap showing the aging trajectories of all differentially expressed genes. We computed each gene's trajectory by averaging z-scaled cell-type-specific trajectories across all cell types. We performed hierarchical clustering on the genes to form six gene modules. B) Left: heatmap showing the GSEA result for neuropeptide signaling (GO:0007218) measured by signed -log10(FDR) between each pair of consecutive time points for non-neuronal cell types; right, heatmap showing the GSEA result for axon guidance (GO:0008045) measured by signed -log10(FDR) between each pair of consecutive time points for non-neuronal cell types.

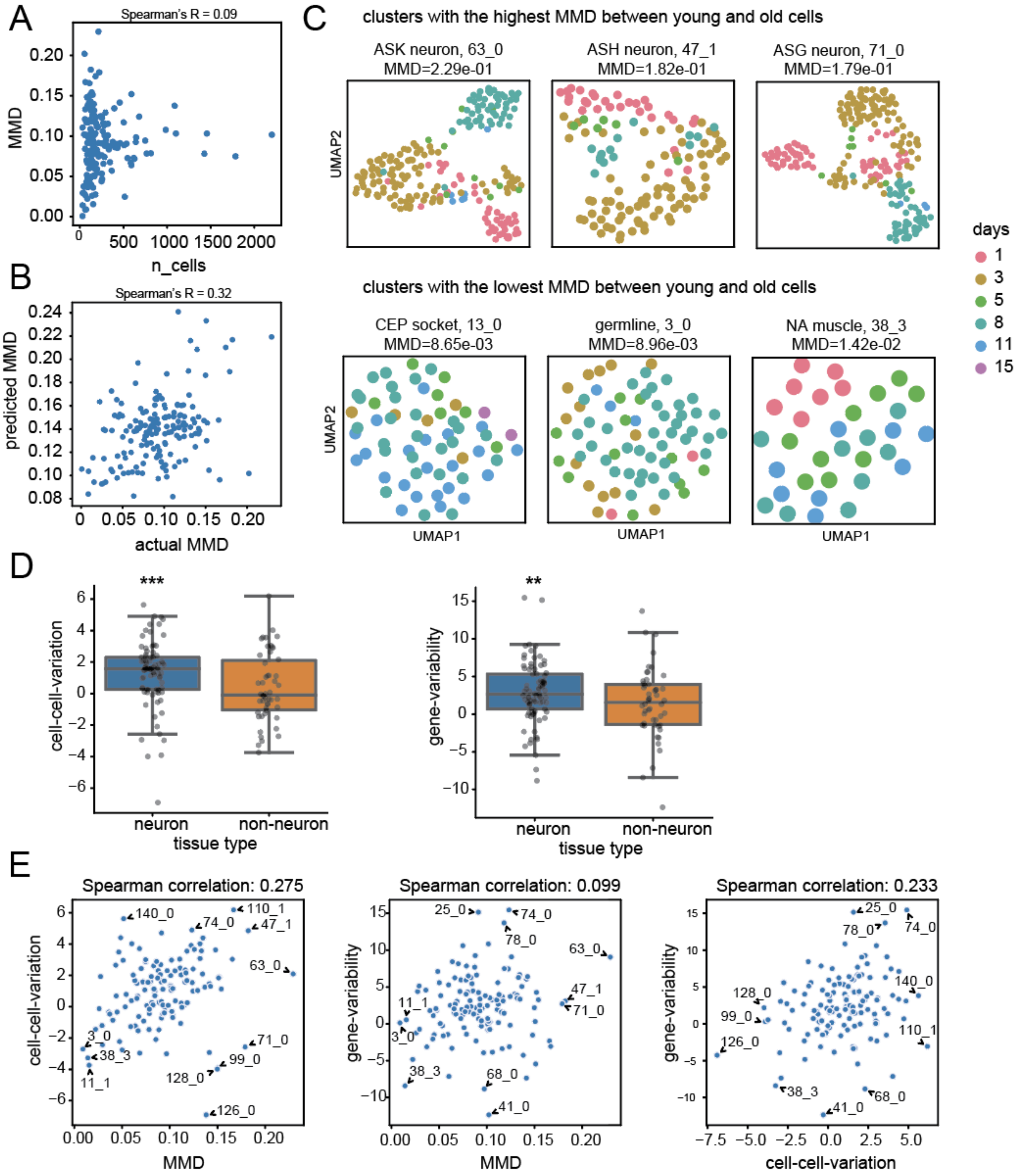


**Figure S5**. Additional comparisons of cell-type global transcriptome changes with age, related to Figure 5.

A) Spearman's correlation of total cell count and estimated MMD distance between young and old cells for each cell type cluster. Cluster cell count does not influence MMD between young and old samples (R = 0.09). B) Spearman's correlation of estimated MMD for each cell type cluster and its closest cell type cluster (based on baseline transcriptome). $p$<2.58e-05. C) Upper panel: UMAP visualization of cells in the clusters 63_0, 47_1 and 71_0 with the highest MMD distance between young and old cells. Lower panel: UMAP visualization of cells in the clusters 13_0, 3_0 and 38_3, whose MMD distance between young and old cells is insignificant. D) Left: Boxplot comparing cell-cell variation and (right) gene-variability for neurons versus other cell types. (*** $p$<1.94e-03, * $p$<2.65e-02). E) Left: Scatterplots for correlation between MMD and cell-cell variation, Middle: correlation between MMD and gene-variability, and Right: correlation between cell-cell variation and gene-variability.

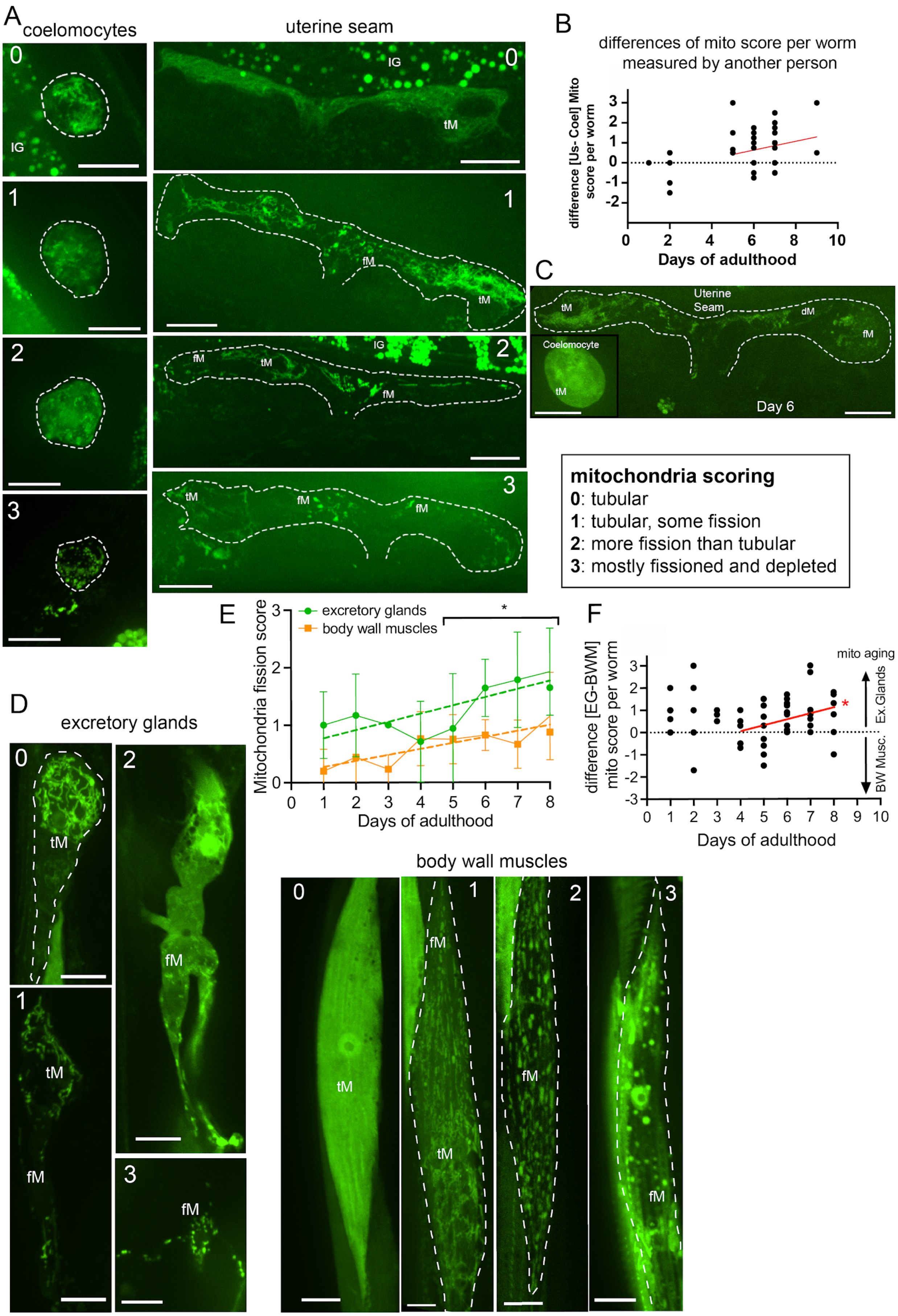

A
coelomocytes
uterine seam
IG
tM
fM
B
differences of mito score per worm measured by another person
difference [Us- Coel] Mito score per worm
Days of adulthood
C
Uterine Seam
dM
Coelomocyte
Day 6
mitochondria scoring
0: tubular
1: tubular, some fission
2: more fission than tubular
3: mostly fissioned and depleted
D
excretory glands
E
excretory glands
body wall muscles
Mitochondria fission score
Days of adulthood
F
difference [EG-BWM] mito score per worm
mito aging
Ex.Glands
BW Musc.
Days of adulthood
body wall muscles

**Figure S6**. Additional quantification of mitochondrial morphology changes with age, related to Figure 6.

A) Left: representative images of the 4 levels of mitochondria fission of coelomocytes ; Right: uterine seam cells as scored. C1/C2: Coelomocyte cells 1 and 2. B) Same as Figure 6C scored by another person. linear regression different from zero *p*=0.038. C) One extra representative image as shown in Figure 6A. D) Left: representative images of the 4 levels of mitochondria fragmentation (named fission in the figure) of excretory glands ; Right: body wall muscles as scored. E) Mitochondria fragmentation average score during aging in excretory glands and body wall muscles. unpaired t test * *p*=0.024. Dashed lines: linear regression. F) Comparisons of excretory glands and body wall muscle mitochondrial fragmentation scores in the same animal during aging. * linear regression significantly different from zero *p*=0.013. dM: depleted mitochondria. fM: fragmented mitochondria. tM: tubular mitochondria. IG: autofluorescent intestinal granules. C1/C2: Coelomocyte 1/2. Scale bar : 10μm.

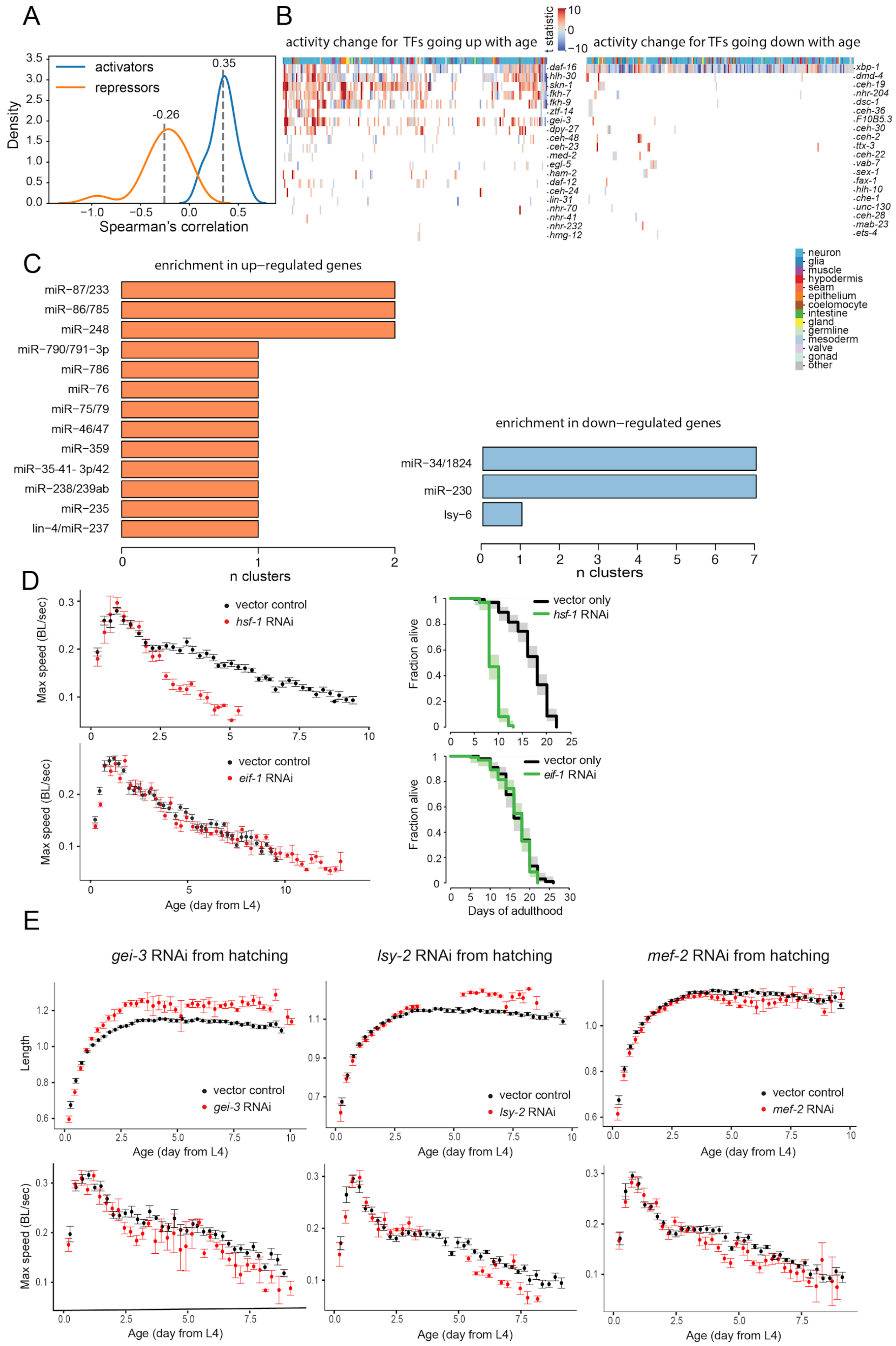
A
activators
repressors
Density
Spearman's correlation
0.35
-0.26
B
activity change for TFs going up with age
t statistic
activity change for TFs going down with age
daf-16
hlh-30
skn-1
fkh-7
fkh-9
ztf-14
gei-3
dpy-27
ceh-48
ceh-23
med-2
egl-5
ham-2
daf-12
ceh-24
lin-31
nhr-70
nhr-41
nhr-232
hmg-12
xbp-1
dmd-4
ceh-19
nhr-204
dsc-1
ceh-36
F10B5.3
ceh-30
ceh-2
ttx-3
ceh-22
vab-7
sex-1
fax-1
hlh-10
che-1
unc-130
ceh-28
mab-23
ets-4
neuron
glia
muscle
hypodermis
seam
epithelium
coelomocyte
intestine
gland
germline
mesoderm
valve
gonad
other
C
enrichment in up–regulated genes
miR–87/233
miR–86/785
miR–248
miR–790/791–3p
miR–786
miR–76
miR–75/79
miR–46/47
miR–359
miR–35-41- 3p/42
miR–238/239ab
miR–235
lin–4/miR–237
n clusters
enrichment in down–regulated genes
miR–34/1824
miR–230
lsy–6
n clusters
D
Max speed (BL/sec)
vector control
hsf-1 RNAi
eif-1 RNAi
Age (day from L4)
Fraction alive
vector only
hsf-1 RNAi
eif-1 RNAi
Days of adulthood
E
gei-3 RNAi from hatching
lsy-2 RNAi from hatching
mef-2 RNAi from hatching
Length
vector control
gei-3 RNAi
lsy-2 RNAi
mef-2 RNAi
Age (day from L4)
Max speed (BL/sec)

**Figure S7**. Additional analysis of transcription factor activity changes with age and their effect on longevity, related to Figure 7.

A) Distribution of Spearman's correlations between changes in expression (log2FC) and changes in target expression (difference in AUCell) for each TF across cell types that they are expressed in. We considered a TF to be a putative activator if the expression-activity correlation across cells (with age regressed out) is > 0.1, and a putative repressor if the expression-activity correlation across cells (with age regressed out) is < -0.1.
B) Left**:** heatmap of differential AUCell score (t statistic) of TF target genes' log2FC during aging for the top 20 TFs with the most positive ; Right: negative average log2FC across cell types that they are expressed in. A blank indicates that the TF is not expressed in that cell type.
C) Left: Barplot of miR families that are significantly enriched in up-regulated genes during aging (padj < 0.01, binomial test). The x-axis indicates how many cell type clusters they are significantly enriched in. Right: Barplot of miR families that are significantly enriched in down-regulated genes during aging (padj < 0.01, binomial test).
D) Healthspan and lifespan of the RNAi knockdown of previously described genes *hsf-1* and *eif-1*. Left panels: Decline in maximum speed (healthspan) after adulthood RNAi treatment. Right panels: Survival after adulthood RNAi treatment.
E) Average worm length (upper panels) and healthspan (lower panels) of the RNAi knockdown of *gei-3*, *lsy-2* and *mef-2* genes initiated from hatching (L1 stage).

#### Supplemental Notes 1, related to Figure 1

##### Germ, embryonic and sperm cells removal

Germ cells vastly outnumber the somatic cells after the initial disassociation (from about 50% up to 91% of cells depending on the age, in *gon-2(q388ts*) mutant). Due to the incomplete penetrance of *gon-2(q388ts)* mutation and the FACS sorted diploid germ cells, we still observed an over-representation of germ cells on the UMAP, as determined by germline marker gene set AUCell scores (Figure SN1A). The same analysis was performed for sperm cells and revealed a small cluster compared to oocytes. In addition, we observed a negligible amount of embryonic cells showing that our final cell population did not contain eggs or early embryos (Figure SN1A).

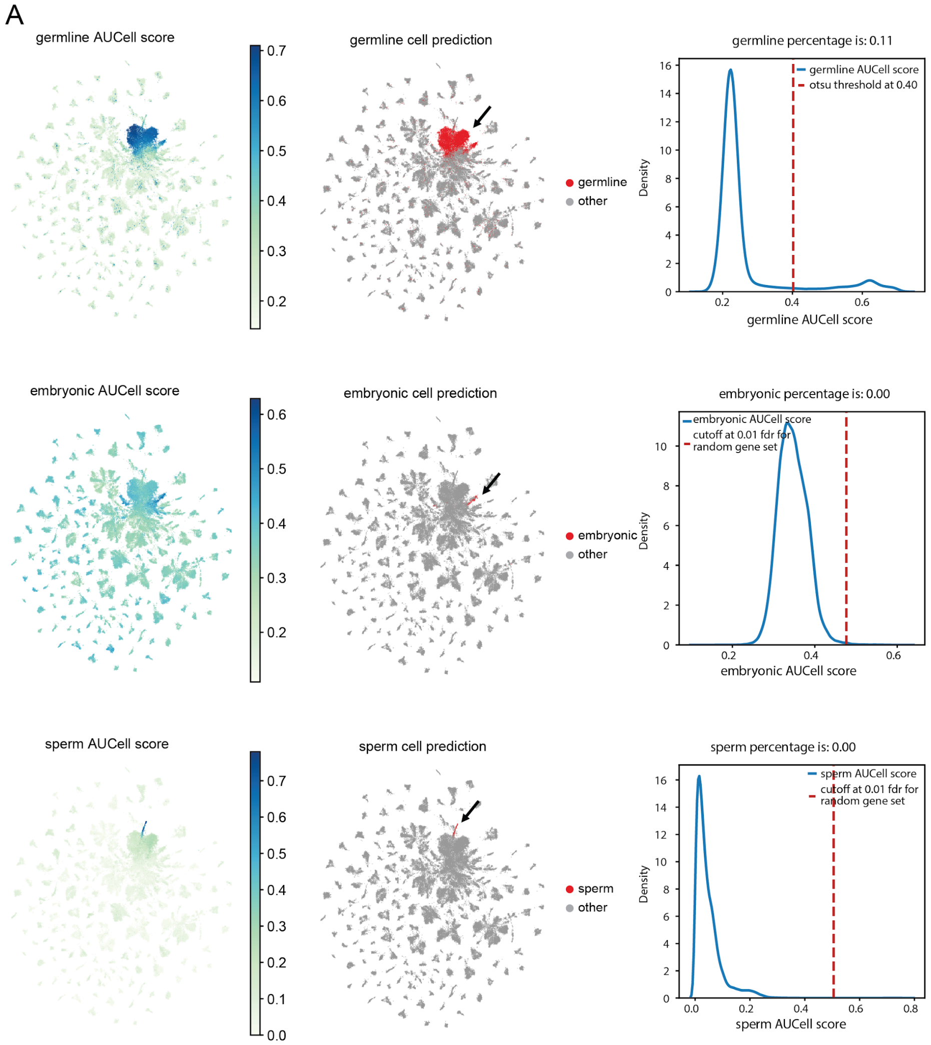


**Figure SN1.** A) First row: (left) UMAP showing germline AUCell score for all cells; (middle) UMAP showing predicted germline cells; (right) distribution of AUCell scores for

all cell*s and the cutoff used to label germline cells. AUCell is a gene set enrichment scoring method, described in more detail in Method Details.*
*Second row: (left) UMAP showing embryonic AUCell score for all cells; (middle) UMAP showing predicted embryonic cells; (right) distribution of AUCell scores for all cells and the cutoff used to label embryonic cells.*
*Third row: (left) UMAP showing sperm AUCell score for all cells; (middle) UMAP showing predicted sperm cells; (right) distribution of AUCell scores for all cells and the cutoff used to label sperm cells.*

**Details on additional ambient RNA removal**

We show *col-140* as an example of residual ambient RNA signal even after CellBender ambient correction and Solo doublet removal (Figure SN1B). We observed that: 1) among most abundant mRNAs in the ambient RNA profile, only cuticle collagen genes are not corrected by CellBender; 2) these residual ambient RNAs are specific to day1 sample;  3) expression of cuticle collagen genes are highly correlated.

We performed additional day1 cell filtering for the ambient RNA. 1) We annotated d1 cell type clusters as “cuticle cluster” or “non-cuticle cluster” based on median *col-140* UMI count > 0 or not. 2) Then we examined the distribution of the cuticle AUCell scores for “cuticle clusters”, and “non-cuticle clusters’, and observed a clear separation (Figure SN1C). 3) We removed all d1 cells in the “non-cuticle cluster” whose cuticle AUCell score > 0.1.

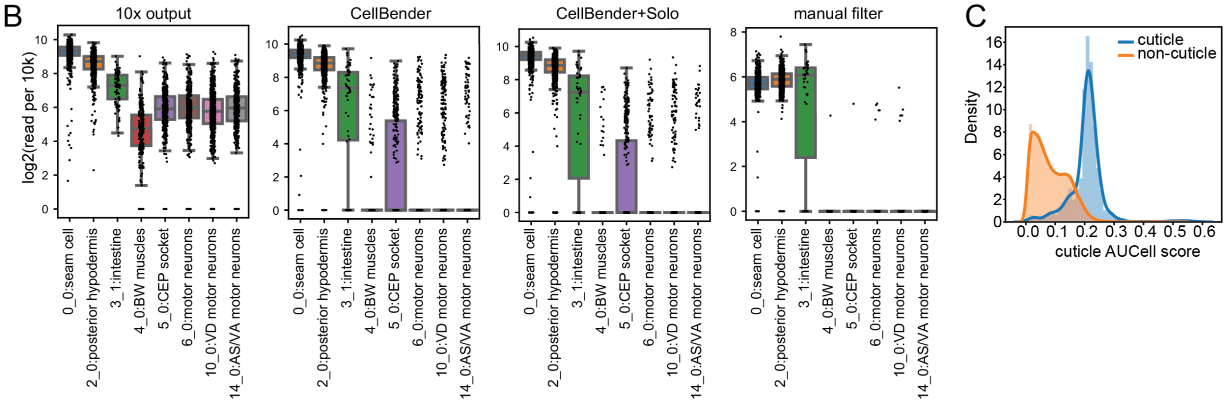


***Figure SN1.*** *B) col-140 UMI count in selected cell type clusters in 10x output, after CellBender correction, after CellBender and Solo filtering, and after additional ambient removal step for the d1 sample. C) distribution of cuticle AUCell scores for cells in cuticle clusters (blue), and cells in non-cuticle clusters (orange).*

#### **Somatic gonad cell-type integrity in *gon-2(q388)***

Because we used a strain carrying *gon-2(q388)* mutation that partially prevents the development of the gonadal tissues, we verified the integrity of these clusters. We compared them (strain CF4596) with those from CF512, a strain carrying a wild-type copy of *gon-2* that was single-cell RNA-sequenced for a preliminary experiment (full data set not published). CF512 carries *fer-15(b26)* and *fem-1(hc17)* alleles that prevent the normal development of their sperm cells. We showed that *gon-2(q388)* did not alter the expression of genes assigned to gonad cells (Figure SN1D). The expression of 4 cell types/clusters including sheath cells, distal tip, vulva uv1 and uv3 showed a high level of correlation with those from CF512.

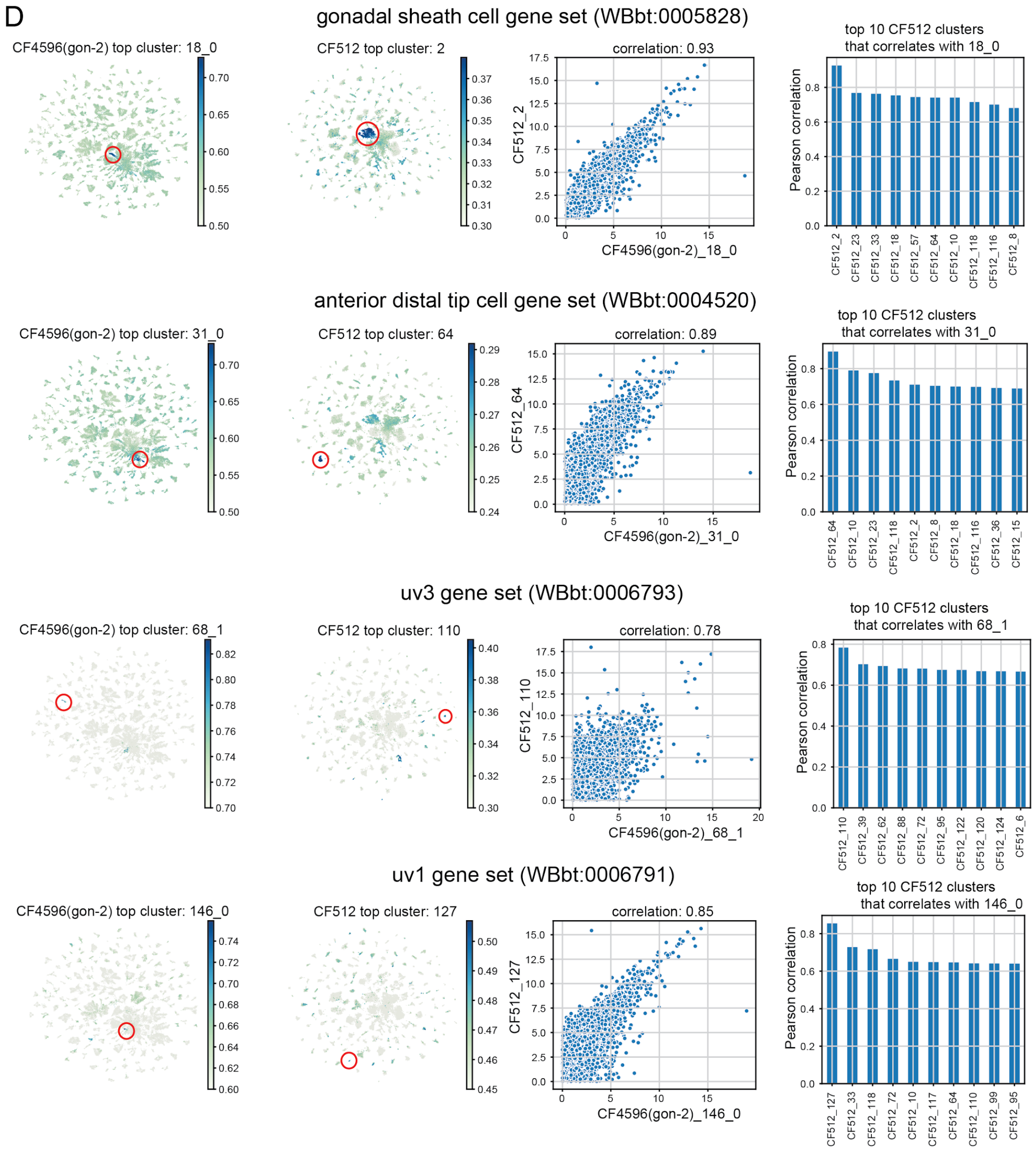

***Figure SN1.*** *D) Cluster-to-cluster comparison of somatic gonad clusters from a gon-2(q388) mutant (CF4596) and a wild type gon-2 (CF512). In CF4596, Cluster 18_0 is annotated as gonadal sheath cell (Wbbt:0005828), cluster 31_0 is annotated as anterior distal tip cell (Wbbt:0004520), cluster 68_1 is annotated as uv3 (Wbbt:0006793), and cluster 146_0 is annotated as uv1 (WBbt:0006791). From left to right, the top row shows: 1) AUCell score for gonadal sheath cell in CF4596. 2) AUCell score for gonadal sheath cell in CF512. Cluster 2 is annotated as the gonadal sheath cell cluster. 3) The correlation of gene expression between the gonadal sheath cell clusters in CF4596 and in CF512. We observe a Pearson correlation of 0.93. 4) Top 10 CF512 cell type clusters that correlate with cluster 18_0 in CF4596. Cluster 2 in CF512 has the highest correlation with 18_0 in q388. The following rows show the same comparison, for different anatomies.*

**Comparison of pre-FACS and post-FACS intestine cluster**

We processed the pre-FACS sample in the same way as the post-FACS sample. We annotated whether each cell is a germ cell based on the AUCell score of the germline marker gene set. We found that > 86% cells from the pre-FACS sample are germ cells. This observation drives us to perform FACS gating to enrich somatic cells.

After filtering out germ cells in the pre-FACS sample, we performed unsupervised Leiden clustering on the sample, and identified the intestine cluster based on expression of key transcription factors *elt-2* and *elt-7*. There are 338 cells in the intestine cluster. Plotting anterior and posterior intestinal markers on these cells, we are able to see cells belonging to anterior intestine and those belonging to posterior intestine are clearly separated by these markers, suggesting good resolution on intestine annotation in this sample.

Next, we evaluate the quality of the annotated intestinal clusters in the post-FACS sample. In the post-FACS sample, cells annotated as intestine cells (cluster 3_1, 3_2, 3_4, 3_5) also show *elt-2* and *elt-7* expression, although at lower level. When we correlate all cell type clusters in the post-FACS sample with the intestine cluster in the pre-FACS sample, we observe that clusters 3_1, 3_2, 3_4, 3_5 correlate best with intestine cluster in pre-FACS sample, suggesting that cells in these clusters do capture transcriptional signatures of intestinal cells. In addition, we compared the marker genes derived from 3_2 with intestinal markers genes from the pre-FACS sample, and observed that 38 of them overlap out of the top 100.

In conclusion, we think that clusters in the post-FACS sample annotated as intestinal cells carry transcriptional signatures of intestinal cells. However, due to the polyploidy of intestinal cells, they might be removed in the FACS filtering process. We thus also provide the intestinal markers derived from the pre-FACS sample for the readers' reference.

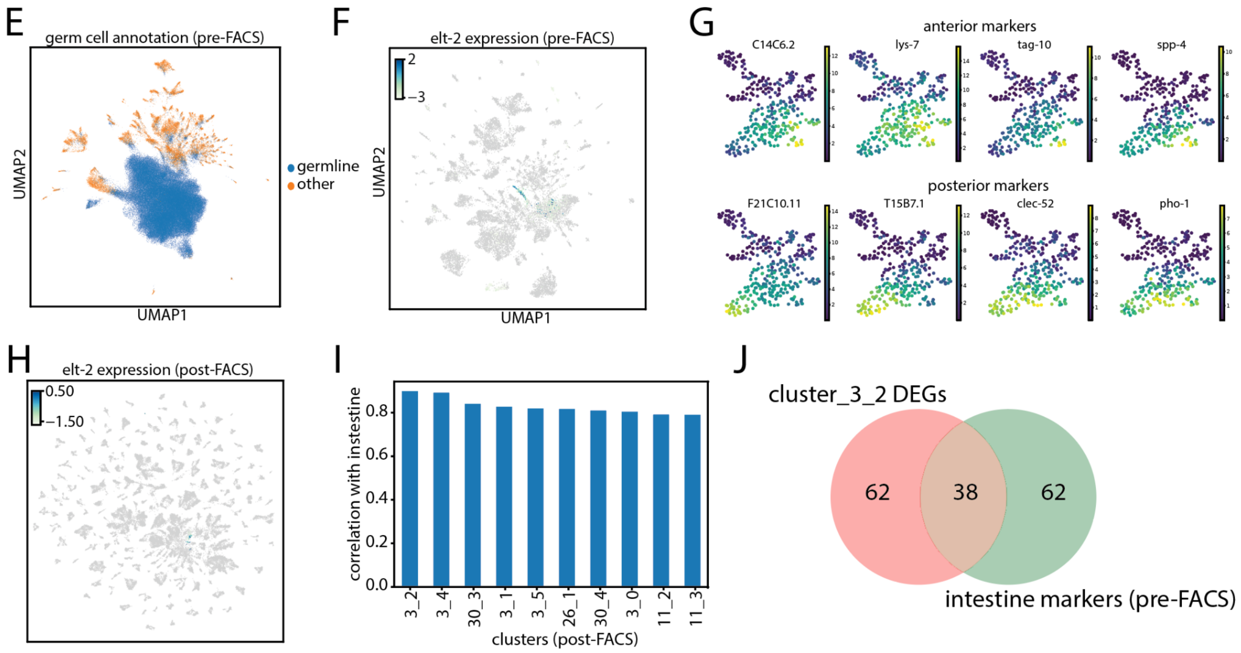


***Figure SN1.*** E) UMAP showing all cells in pre-FACS sample. Cells are colored by whether it is a germ cell or not. F) UMAP after filtering out germ cells. We colored cells by denoised *elt-2* expression level. Denoised expressions are normalized to log2(read per 10k). G) UMAP showing expression of anterior and posterior intestinal markers. Denoised expressions are normalized to log2(read per 10k). H) *elt-2* expression for cells in post-FACS sample (log2 read per 10k). I) Spearman correlation of the aggregated transcriptional profile of each cell type cluster in post-FACS sample with the intestinal cluster in the pre-FACS sample. J) overlap of markers derived from cluster 3_2 in the post-FACS sample, and the intestinal markers from the pre-FACS sample.

#### Supplemental Notes 2, related to Figure 3

##### Specific analysis of cell types aging signatures

To perform a gene set enrichment analysis (GSEA) of GO terms with age, we tested 2,166 *C. elegans* GO terms and found 279 of them associated with age in at least one cell type cluster (q-value < 0.01). We removed redundant GO terms by hierarchical clustering (Figure SN2A). In the main manuscript we present the top 20 most changing

GO terms that are fully detailed here (Figure SN2B-C). The full list of the top 100 GO terms with significant changes during aging are presented here (Figure SN2D). We observed very few GO terms changing consistently in all clusters during aging. Inversely we found that most GO term changes with age occur in a cell type- or tissue-specific fashion. It was the case for two types of extracellular matrix gene sets (collagen and basement membrane genes, Figure SN2D-F) as well as for proteasome genes Figure SN2G).

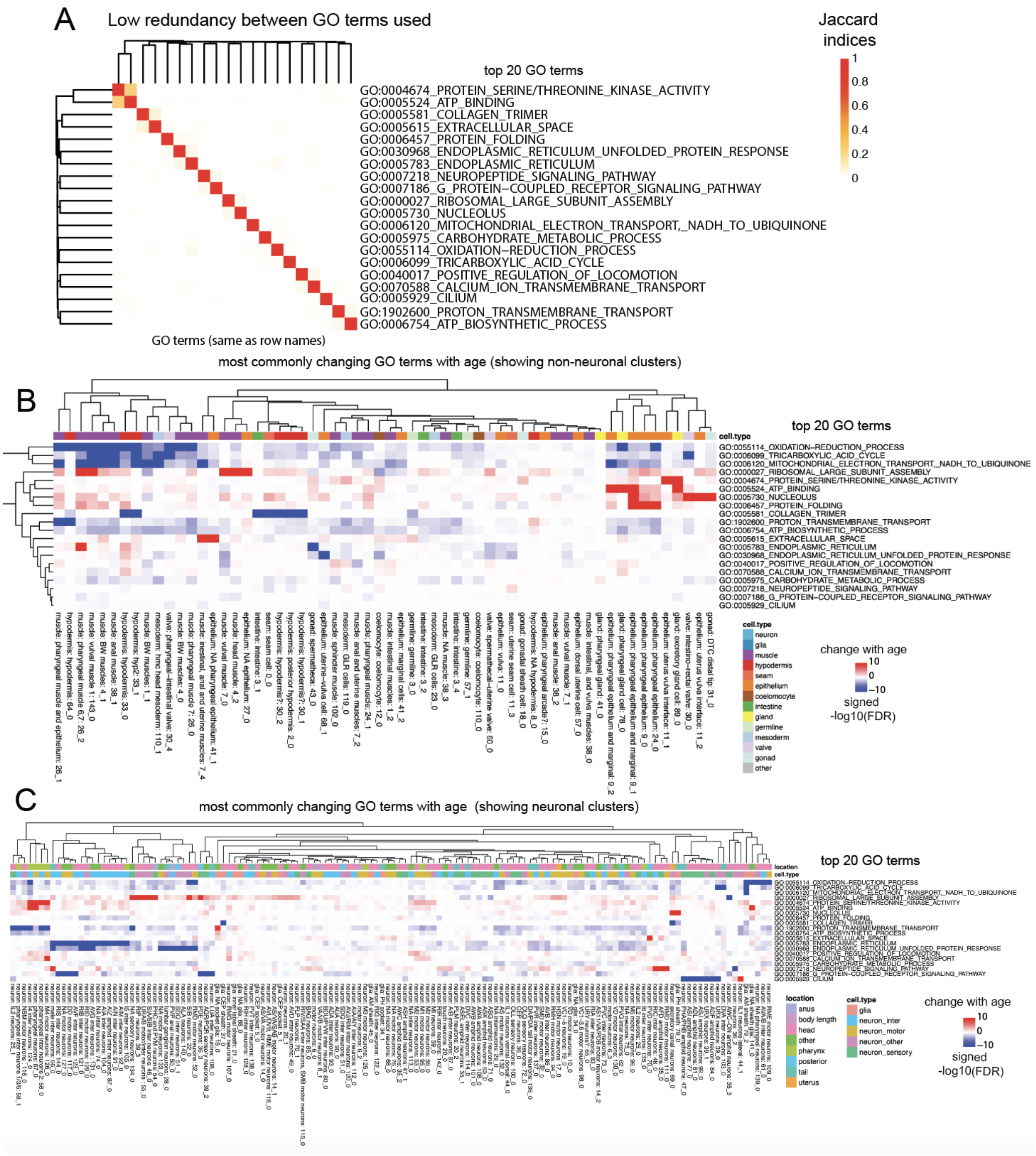


*Figure SN2. A) Heatmap of Jaccard indices between every pair of GO terms for top 20 GO terms with most changes (same GO terms as Fig. 3C). B) GSEA results in non-neuronal cell types for 20 representative GO terms with significant changes in most clusters. Rows are the GO terms, and columns are the clusters summarized (averaged) to a high-level tissue annotation. Color indicates the signed -log10(FDR) of the GSEA results. Positive indicates that the gene set is enriched in old cells, whereas negative*

*indicates that the gene set is enriched in young cells. C) Same as B) for neuronal clusters.*

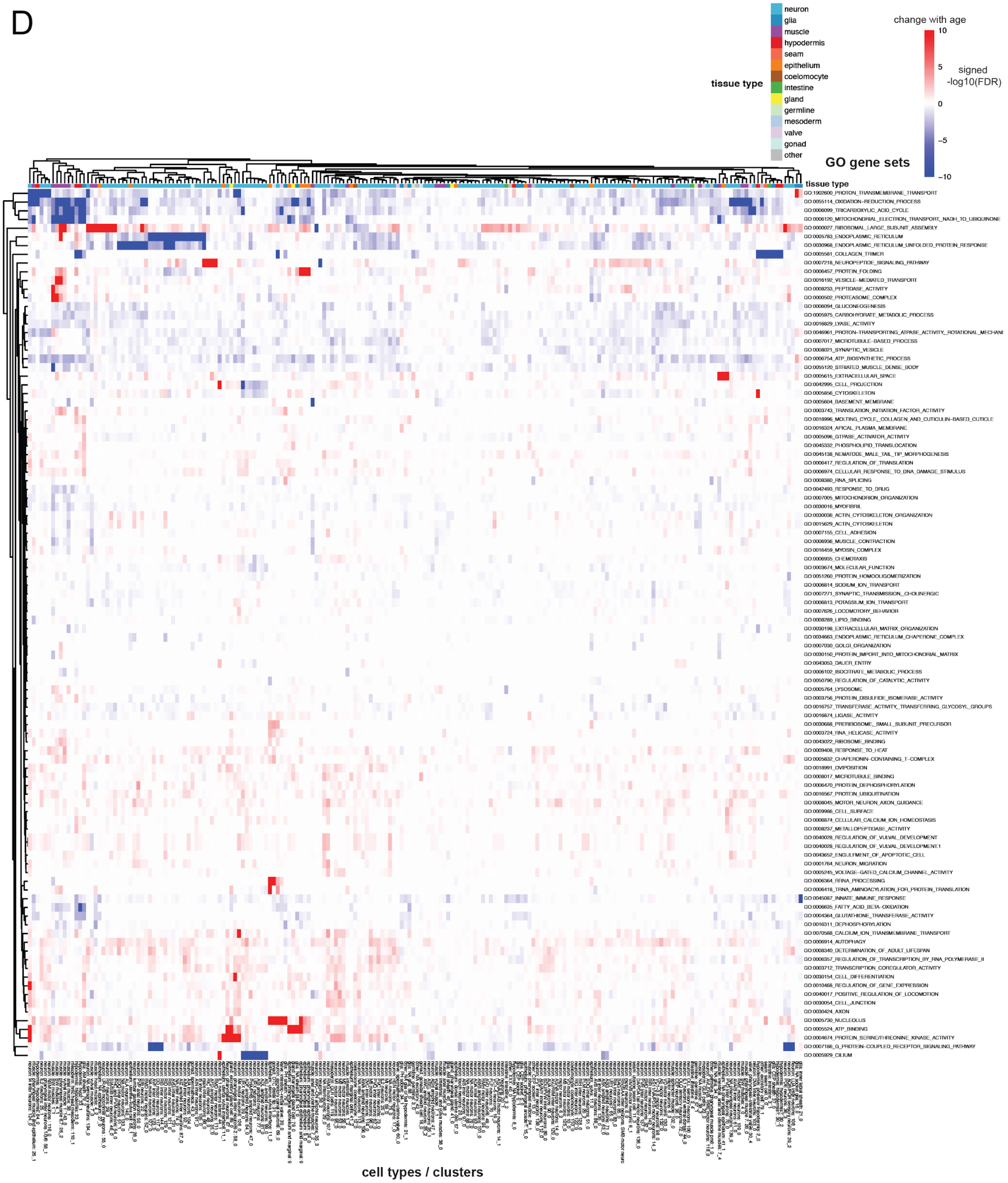


*Figure SN2. D) GSEA results for the top 100 non-redundant GO terms with significant changes during aging. Rows are the GO terms, and columns are the clusters. Color indicates the signed -log10(FDR) of the GSEA results. Positive indicates that the gene*

*set is enriched in old cells, whereas negative indicates that the gene set is enriched in young cells.*

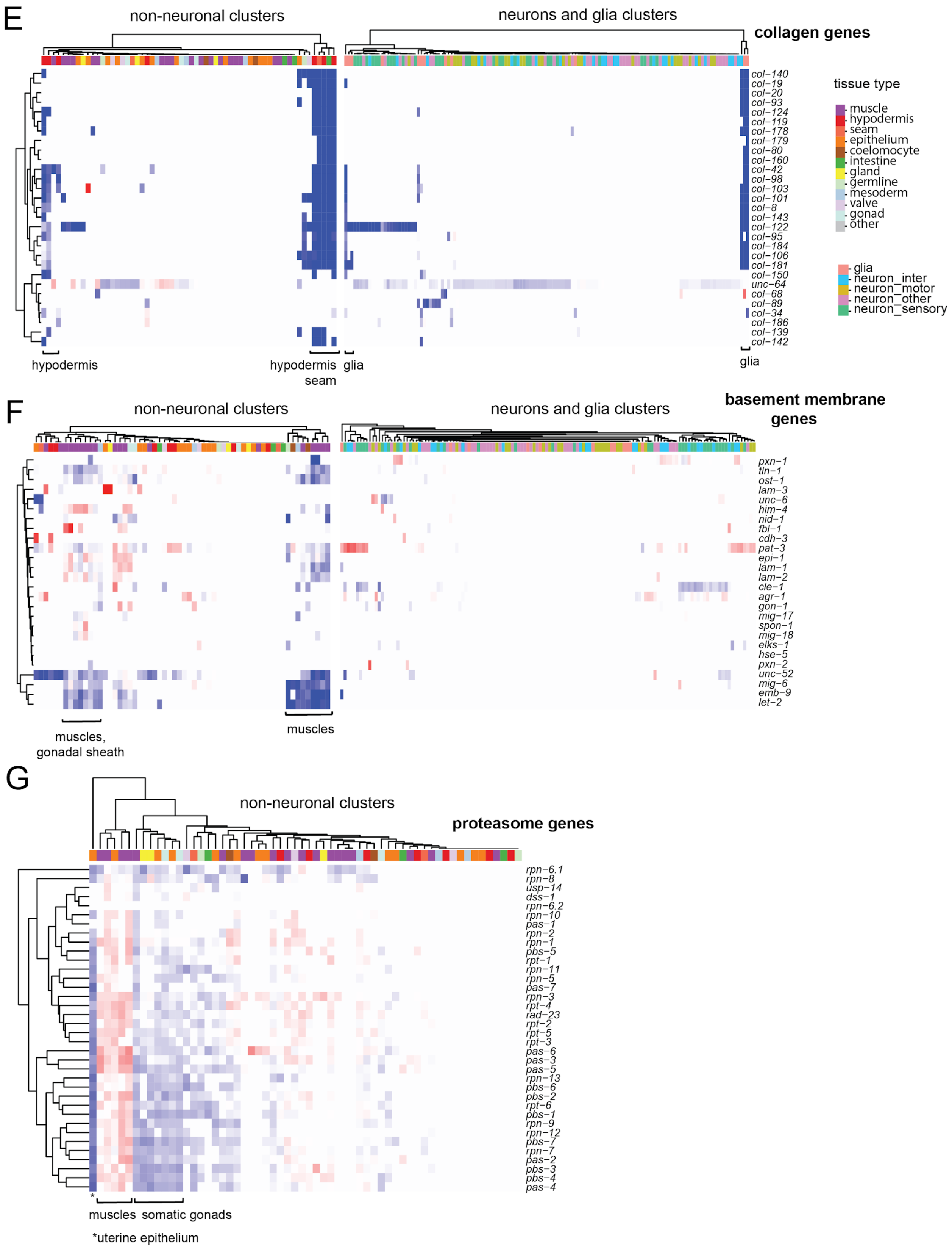
E
non-neuronal clusters
neurons and glia clusters
collagen genes
col−140
col−19
col−20
col−93
col−124
col−119
col−178
col−179
col−80
col−160
col−42
col−98
col−103
col−101
col−8
col−143
col−122
col−95
col−184
col−106
col−181
col−150
unc−64
col−68
col−89
col−34
col−186
col−139
col−142
hypodermis
hypodermis
seam
glia
glia
tissue type
muscle
hypodermis
seam
epithelium
coelomocyte
intestine
gland
germline
mesoderm
valve
gonad
other
glia
neuron_inter
neuron_motor
neuron_other
neuron_sensory
F
non-neuronal clusters
neurons and glia clusters
basement membrane genes
pxn−1
tln−1
ost−1
lam−3
unc−6
him−4
nid−1
fbl−1
cdh−3
pat−3
epi−1
lam−1
lam−2
cle−1
agr−1
gon−1
mig−17
spon−1
mig−18
elks−1
hse−5
pxn−2
unc−52
mig−6
emb−9
let−2
muscles, gonadal sheath
muscles
G
non-neuronal clusters
proteasome genes
rpn−6.1
rpn−8
usp−14
dss−1
rpn−6.2
rpn−10
pas−1
rpn−2
rpn−1
pbs−5
rpt−1
rpn−11
rpn−5
pas−7
rpn−3
rpt−4
rad−23
rpt−2
rpt−5
rpt−3
pas−6
pas−3
pas−5
rpn−13
pbs−6
pbs−2
rpt−6
pbs−1
rpn−9
rpn−12
pbs−7
rpn−7
pas−2
pbs−3
pbs−4
pas−4
*
muscles
somatic gonads
*uterine epithelium

*Figure SN2. E) Heatmap of gene expression log2FC in old versus young cells for collagen genes GO term (GO:000005581) (rows) for all non-neuronal cell types clusters (columns). F) Heatmap of gene expression log2FC in old versus young cells for basement membrane genes GO term (GO:000005604) (rows) for all non-neuronal cell types clusters (columns). G) Heatmap of gene expression log2FC in old versus young cells for proteasome genes GO term (GO:00000502) (rows) for all non-neuronal cell types clusters (columns).*

#### Supplemental Notes 3, related to Method Details

The figures below support the protocol of cell isolation and cell sorting. Figure SN3A shows how to make a custom 35µm nylon filter (Kaletsky et al. 2016) from a commercial cell strainer (Falcon ref352340). Pictures in SN3B show the cells in solution before they are processed through the FACS sorter. The graph in Figure SN3C shows the gating we used to enrich 2N ploidy cells and remove most germ lines cells.

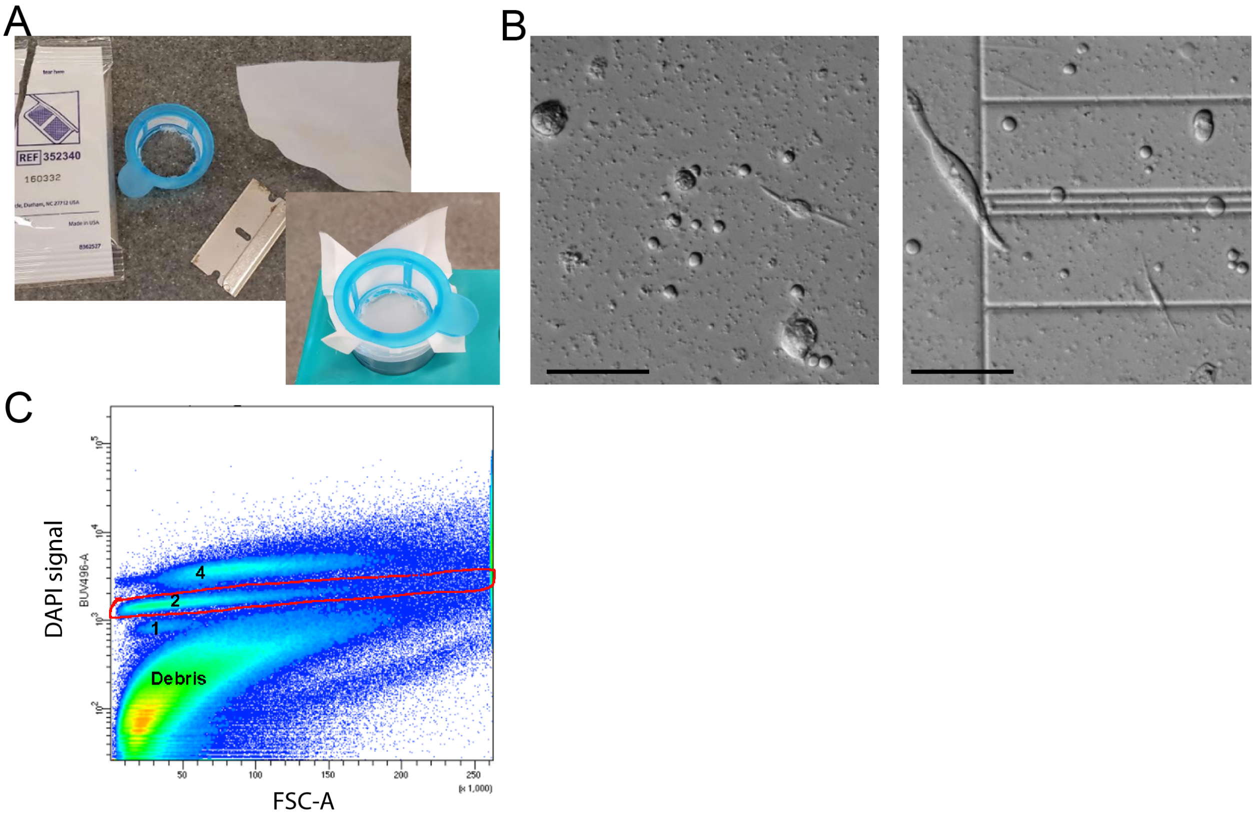


*Figure SN3. Adult C. elegans single-cell isolation.*
*A) Setup to adapt a 35µm nylon mesh filter on a commercial strainer.*

*B) DIC pictures of cells in solution in the hemocytometer before they are flowed through the cell sorter. Scale bar is 0.5μm.*
*C) Representation of sorting events and gating strategy. A DAPI-compatible filter versus forward scatter graph was used to gate. The red line represents the gating strategy used to enrich 2N somatic cells. 1: 1N-DNA cells (this population is visible only in younger worms); 2: 2N-DNA cells; 4: 4N-DNA cells.*
